# Skeletal Muscle Regeneration is Accelerated Following Injection of Time Release Ion Matrix in Injured Mice

**DOI:** 10.1101/2025.11.21.689759

**Authors:** Jacob A. Kendra, Jiwei Hao, Dillon R. Harris, Rebekah L. Blatt, Alexandra G. Naman, Seth D. Goble, Lydia D. Garcia, Richard K. Brow, Aaron B. Morton

## Abstract

**Introduction:** Skeletal muscle injury remains a significant cause of disability with limited treatment options. Here we report accelerated skeletal muscle regeneration following injection of an inorganic biomaterial alone, cobalt oxide time-release ion matrix (CoO-TRIM).

**Methods:** The tibialis anterior (TA) muscle of adult C57BL/6J mice was injured with 70 µL of barium chloride, with uninjured limbs serving as contralateral controls. Following the acute inflammatory phase (3 days post injury, dpi), mice were randomly separated into three groups: untreated controls, 70 µl sterile saline injection alone (vehicle control) or 70 µl CoO-TRIM (5 µg/µl) and evaluated at 8- and 14 dpi.

**Results:** 14 dpi, injured TA muscles receiving a single injection of CoO-TRIM exhibited greater recovery of maximal force production (means ± SEM: Healthy; TRIM, 31.8 ± 0.6 N/cm^3^; Untreated, 31.7 ± 0.8 N/cm^3^; Saline, 31.6 ± 0.8 N/cm^3^, Injured; TRIM, 31.8 ± 0.8 N/cm^3^; Untreated, 26.1 ± 1.1 N/cm^3^, P= <0.01 *vs.* TRIM Injured; Saline, 26.0 ± 1.3 N/cm^3^, P= <0.01 *vs.* TRIM Injured), increased fiber size (Healthy; TRIM, 2466 ± 128 µm^2^ *vs.* Untreated, 2351 ± 131 µm^2^; *vs.* Saline, 2460 ± 129 µm^2^, Injured; TRIM, 2179 ± 63 µm^2^ *vs.* Untreated, 1525 ± 52 µm^2^, P= <0.01; *vs.* Saline, 1470 ± 99 µm^2^, P= <0.01), and accelerated muscle regeneration (TRIM, 32.7 ± 6.8 eMyHC^+^ fibers/mm^2^ *vs.* Untreated, 91.8 ± 11.8 eMyHC^+^ fibers/mm^2^, P= <0.01; *vs.* Saline, 94.0 ± 9.6 eMyHC^+^ fibers/mm^2^, P= <0.01). Vascular endothelial growth factor was elevated 14 dpi (TRIM, 17.3 ± 2.9 pg/mg *vs.* Untreated, 10.3 ± 1.1 pg/mg, P= 0.04; Saline, 5.5 ± 1.2 pg/mg, P= <0.01), with increased muscle microvascular area (TRIM, 119.9 ± 4.2 µm^2^/fiber *vs.* Untreated, 94.9 ± 4.2 µm^2^/fiber, P= <0.01; Saline, 91.3 ± 4.0 µm^2^/fiber, P= <0.01) following CoO-TRIM treatment. There were early increases in inflammatory responses 8 dpi in injured TA muscles receiving CoO-TRIM (IL-6; TRIM, 29.1 ± 8.2 pg/ml *vs.* Untreated, 12.4 ± 1.5 pg/ml, P= 0.02; Saline, 11.7 ± 0.8 pg/mg, P= 0.01), with early resolution of degenerative inflammatory cytokines and elevated regenerative cytokines 14 dpi (IL-10; SEM: TRIM, 2.4 ± 0.1 pg/ml *vs.* Untreated, 2.1 ± 0.1 pg/ml, P= 0.25; Saline, 2.0 ± 0.1 pg/ml, P= 0.04). No differences were observed in healthy contralateral limbs following treatment with CoO-TRIM compared to healthy control mice at 8- and 14 dpi.

**Discussion:** This work suggests CoO-TRIM enhances muscle regeneration following injury, possibly through an immune cell-mediated mechanism.

**Statement of Clinical Relevance:** This is the first report of an injected, inorganic biomaterial alone to accelerate regeneration of injured skeletal muscle with no observed effects on healthy muscle. These findings suggest enhancements to skeletal muscle regeneration following CoO-TRIM treatment may be immune cell mediated with an increased local inflammatory response and subsequent improvements to muscle microvasculature and myogenic regulatory factors.

## Introduction

Skeletal muscle injuries are the leading cause of disability for warfighters and represent nearly 50% of all sports-associated injuries(1, 2). The United States Department of Defense estimates approximately 1.6 million musculoskeletal injuries occur annually with the leading cause of injury classified as overuse-induced damage from sport and physical training(3). Regeneration in skeletal muscle is tightly regulated through biophysical and biochemical cues, removing tissue debris prior to revascularization and reformation of contracting myofibrils(4). Following injury to otherwise healthy muscle, resident neutrophils and macrophages initiate a proinflammatory response, recruiting circulating macrophages to the muscle microenvironment, elevating levels of the proinflammatory cytokines tumor necrosis factor-alpha (TNF-α), and interleukins (IL) 1-β, IL-6, to facilitate the removal of damaged myofilaments and organelles at the site of injury(5). Proinflammatory macrophages transition towards pro-regenerative macrophages, both secreting cytokines and growth factors, i.e., TNF-α, IL-1β, IL-6, and IL-10, vascular endothelial growth factor (VEGF) and insulin growth factor 1 (IGF-1). These biochemical cues are required for angiogenesis and satellite cell proliferation, prompting functional revascularization, maturation, and remodeling of newly formed fibers(6–8). While the regenerative events in skeletal muscle have been widely investigated, no current standard treatment exists to enhance skeletal muscle regeneration following injury.

Tissue engineering and biomaterials have emerged as promising strategies to enhance skeletal muscle regeneration. Organic scaffolds (i.e., hydrogels, collagen, fibrin, decellularized extracellular matrix) provide bioactive cues resembling native muscle extracellular matrix, mimicking the natural regeneration cascade, while synthetic and hybrid polymers possess tunable mechanical properties and controlled degradation rates creating effective delivery vehicles(9, 10). Despite these advances, cells and/or growth factors are frequently incorporated to achieve functional vascularization and activation of satellite cells to promote skeletal muscle regeneration(11–13). While transplantation of freshly isolated satellite cells has improved muscle repair through engraftment, proliferation and survival, these advancements still require cell removal from the satellite cell niche contributing to loss of stemness(14–17). Bolus deliveries of VEGF and IGF-1 enhance the normal regenerative process in skeletal muscle, however limitations remain, in particular, short half-life and loss of bioactivity. When VEGF and IGF-1 were incorporated into biomaterial delivery vehicles, significant improvements to muscle regeneration, vascularization and function were observed(18–21), but the inclusion of such growth factors can be time consuming and financially costly(22).

Time-release Ion Matrices (TRIMs) are an advancing class of inorganic biomaterials capable of regenerating hard and soft tissue through temporal release of ions stimulating active gene expression(23, 24). The original 45S5 silicate-based glass composition, Bioglass^®^, remains the gold standard for biocompatible ceramics, with its composition designed to form a hydroxyapatite layer integrating scaffold to bone(25). However, limitations exist surrounding the use of 45S5 in soft tissue due to several intrinsic material properties. The silicate-based 45S5 composition leads to a rapid ion exchange and alkalization upon dissolution altering the local pH(26). Furthermore, incomplete scaffold degradation is incompatible with the mechanical compliance necessary for integration in dynamic soft tissue such as skeletal muscle(27). These limitations result in poor cell infiltration, limited extracellular matrix remodeling, and inefficient stimulation of satellite cells. Given advances in the design of TRIMs, modified compositions composed of borate and phosphate (borophosphate) have gained interest due to their neutral pH, controlled degradation, resistance to hydroxyapatite formation, and incorporation of therapeutic ions suited for skeletal muscle regeneration(28, 29). While previous work has demonstrated borophosphate TRIMs support angiogenesis and satellite cell activation for enhanced *in vivo* regeneration of volumetric muscle loss (VML) and muscle disease(30), no comprehensive study has investigated the effect of TRIMs on skeletal muscle function, rate of regeneration, or immune cell activity following myotoxin-induced injury.

Here, we investigate skeletal muscle regeneration following the introduction of an inorganic biomaterial, Cobalt Oxide (CoO)-TRIM, to evaluate the regenerative signaling cascade. Using a barium chloride (BaCl_2_) injury model, we demonstrate enhanced functional muscle recovery, increased muscle fiber size, and accelerated regeneration up to 14 days post injury (dpi) following treatment with CoO-TRIM. Interestingly, while we observed no effect of CoO-TRIM in healthy muscle, it significantly augmented immune responses in the injured muscle, with subsequent increases in regenerative growth factor expression and revascularization. These findings suggest CoO-TRIM as a promising approach to support efficient recovery of skeletal muscle structure and function following injury.

## Methods

Detailed methods can be found within the electronic data supplement.

### Biomaterial Fabrication

CoO-TRIM is a CoO-incorporated Na-Ca-borophosphate material with an analyzed composition %wt of: 34.6% B_2_O_3_, 35.3% P_2_O_5_, 14.0% CaO, 12.3% Na_2_O, 3.8% CoO. CoO-TRIM was prepared from mixtures of reagent grade NaPO_3_ (Shanghai Muhong Industrial Co., Ltd., Optical Grade), Ca(PO_3_)_2_ (Shanghai Muhong Industrial Co., Ltd., Optical Grade), H_3_BO_3_ (Fisher, ≥99.5%), Na_2_CO_3_ (Alfa Aesar, 99.5%), CaCO_3_ (Fisher, >98.0%) and CoO (Sigma Aldrich, 99.5%) as raw materials using conventional glass processing methods.

### Ethical approval

Protocols in this study were approved by Animal Care and Use Committees at Texas A&M University (AUP 2022-0215). Animal care was in accordance with the National Research Council’s Guide for the Care and Use of Laboratory Animals (*Eighth Edition, 2011*)

### Animals

Male and Female C57BL/6J mice (RRID: IMSR_JAX:013141, Strain 013141, Jackson Labs; Bar Harbor, ME, USA) mice were housed in the Texas A&M University animal care facilities and studied at 4-5 months of age. Mice were housed on a 12h: 12h light: dark cycle ∼23°C, with fresh water and food available *ad libitum*.

Mice were randomly assigned to one of three experimental groups: 1) Untreated; 2) Saline treated (vehicle control); 3) CoO-TRIM treated. To induce muscle injury *in vivo*, mice were anesthetized via isoflurane inhalation (4% induction, 2% maintenance), the skin was shaved over the anterior hindlimbs, and 1.2% BaCl_2_ (70 uL) was injected unilaterally into either the right or left TA with the uninjured limb serving as a contralateral control. Mice were kept warm during recovery and then returned to their cage followed by 3 days of observation. 3 days post injury (dpi), mice in the Saline and CoO-TRIM experimental groups were anesthetized (as above) and 70 uL of saline or 350 ug of CoO-TRIM suspended in 70 uL of saline (5 ug/ul) was injected into the TA muscle of both hindlimbs. At 8- and 14 dpi, both TA muscles were prepared for *in situ* measurements of maximal isometric force as the criterion for muscle function. The mouse was then euthanized via anesthetic overdose and cervical dislocation. TA muscles from both hindlimbs were removed and stored at −80°C for histological and biochemical analyses.

### Muscle Force

The contractile properties of the tibialis anterior (TA) muscles were evaluated *in situ* in anesthetized mice (1301A 3-in-1 Whole Animal System for Mice, Aurora Scientific, Ontario, Canada). Mice were anesthetized (as above) and weighed. Maximum tetanic contractions were obtained at 120 Hz for 500 ms. Data for force production was acquired using manufacturer software (610A Dynamic Muscle Control/Analysis, Aurora Scientific, Ontario, Canada) on a personal computer. Following contractions, the TA muscles were removed, blotted of excess moisture, and weighed (ME54E, Mettler Toledo; Columbus, OH, USA).

### Immunofluorescence Analysis

Primary antibodies used were mouse anti-embryonic myosin heavy chain (1:5, RRID: AB_528358, Cat. # F1.652, Developmental Studies Hybridoma Bank, The University of Iowa Department of Biology; Iowa City, IA, USA), rat anti-CD31 (1:200, RRID: AB_393571, BD Biosciences, Franklin Lakes, NJ, USA), and rabbit anti-laminin (1:400, Cat.# NC1732938, Fisher Scientific; Hampton, NJ, USA). Secondary antibodies were all from Fisher Scientific (Hampton, NJ, USA): Alexa Fluor 488 Goat anti-mouse (1:400, RRID: AB_2633275, Cat.# PIA32723), Alexa Fluor 647 Goat anti-rat (1:400, RRID: AB_2895299, Cat.# A48265), Goat anti-rabbit Rhodamine (TRITC) (1:400, RRID: AB_90296, Cat.# AP132RMI).

For embryonic myosin heavy chain (eMyHC^+^) analysis, 10x (N.A., 0.40) tile square images were acquired and merged in Leica LAS_X software (RRID: SCR_013673, Leica Microsystems, Deer Park, IL, USA). Using NIH ImageJ (RRID: SCR_003070, NIH, Bethesda, MD, USA), the total number of eMyHC^+^ fibers were counted and normalized to the total area (mm^2^) of the muscle cross section as an index of the rate of regeneration. The total number of regenerated muscle fibers identified by central nuclei were counted in cross section and normalized to the total area (mm^2^) of the muscle cross section as an index of muscle fiber regeneration. For muscle fiber cross-sectional area (CSA) and CD31^+^ analysis, 4-6 randomized 580 x 580 µm regions of interest (ROI) spanning the entire muscle were imaged with a 20x (N.A., 0.75) objective and values across all regions were averaged per muscle. An average of 400 muscle fibers per TA muscle section were evaluated. Muscle fiber CSA was analyzed via semi-automatic muscle segmentation analysis in *NIS*-Elements Advanced Research software (RRID: SCR_014329, Nikon, Melville, NY, USA) presented as µm^2^. To evaluate CSA in the injury groups, only eMyHC^+^ or central nucleated fibers were analyzed to assess CSA of regenerated muscle fibers compared to healthy control group CSA. Fibers were classified as small fibers (<800 µm^2^) or large fibers (>2000 µm^2^). The total area of CD31^+^ staining per fiber was measured to evaluate microvascular area using Aivia machine-learning software (Leica LAS_X Software imported calibration: 0.57 µm/px) (Aivia, Bellevue, WA, USA) and normalized to the total number of muscle fibers in the ROI’s stated above.

### Muscle Tissue Preparation

TA muscle samples frozen for biochemical analyses were homogenized in lysis buffer (pH = 7.4) containing 5 mM Tris-HCI, 5 mM EDTA with protease and phosphatase inhibitors (Sigma-Aldrich, St. Louis, MO, USA). Homogenates were centrifuged at 1,500 *g* for 10 minutes at 4 °C. The supernatant (soluble fraction) was aspirated, and its protein concentration assessed using the Bradford method (Sigma-Aldrich, St. Louis, MO, USA).

### Western Blot Analysis

Protein concentration of each sample was normalized in 4x Laemmli sample buffer (Cat. # 1610747, Bio-Rad, Hercules, CA, USA) containing 5% dithiothreitol. Samples were loaded on 4-20% gradient Criterion TGX Midi-Protein gels (Cat. # 5678095, Bio-Rad, Hercules, CA, USA) for electrophoresis and transferred to LF-PVDF membranes (Cat. # IPFL00010, Millipore, Burlington, MA, USA). Following transfer, membranes were blocked in either 5% milk solution or Intercept TBS Blocking Buffer (Li-Cor Biotechnology, Lincoln, NE, USA) for 1 h at RT; followed by incubation with primary antibodies overnight at 4 °C. Membranes were exposed to either IRDye 680RD Goat anti-Mouse IgG (RRID: AB_10956588, Li-Cor Biotechnology, Lincoln, NE, USA) or IRDye 800CW Goat anti-Rabbit IgG secondary antibodies (RRID: AB_621843, Li-Cor Biotechnology, Lincoln, NE, USA) for 1 h at RT. Western blots were normalized to total protein according to manufacturer recommendations for fluorescent western blotting using Revert Total Protein Stain (Cat. # 92611021, Li-Cor Biotechnology, Lincoln, NE, USA). Individual data points are displayed as fold change (FC) of corresponding Healthy Saline means. Primary antibodies of interest, membranes and total protein stain images are included in the data supplement.

### ELISA

Samples of TA muscle supernatant (50 µL for VEGFA, 100 µL for IGF-1; referred to above) were analyzed using a quantikine enzyme-linked immunosorbent assay (ELISA) to determine VEGFA (RRID: AB_2847842, Cat.# MMV00, R&D Systems, Minneapolis, MN, USA) and IGF-1 (Cat.# ELM-IGF1, RayBiotech, Peachtree Corners, GA, USA) protein concentrations according to the manufacturer’s instructions. TA muscle supernatant (50 µL sample; referred to above) were analyzed to determine cytokine and chemokine concentrations using V-Plex Proinflammatory Panel 1 (mouse) kits (Cat.# K15048D-1, Meso Scale Discovery, Rockville, MD, USA) according to manufacturer’s instructions for the following analytes: IFN-y, IL-1β, IL-2, IL-4, IL-5, IL-6, IL-10, IL-12p70, KC/GRO (CXCL1), TNF-α.

### RNA Isolation and Quantitative RT-PCR

Oligonucleotide gene array (Thermo Fisher Scientific, Waltham, MA, USA) used was *Hif1α* (Mm00468869_m1). For mRNA analysis, housekeeping gene *GAPDH* (Mm99999915_g1) was used. Results are represented as relative fold change in gene expression normalized to the Healthy Untreated controls. For statistics, the relative delta-delta Ct method was used.

### Statistical Analysis

Summary data are reported as means ± standard error mean (SEM). Statistical analyses were performed using Prism 9 software (RRID: SCR_002798, GraphPad Software, La Jolla, CA, USA). Data were analyzed using a two-way ANOVA (Condition x Treatment) with Sidak’s multiple comparisons when appropriate. Values for “*n*” refer to the number of mice studied in each experimental group. *P* ≤ 0.05 was considered statistically significant. A power analysis was conducted *a priori* indicating a minimum n=6 was sufficient to reach a power of 0.8. Additional assays were checked for power *post hoc*.

## Results

### TA Muscle Force is Increased Following CoO-TRIM Treatment

No changes were observed in maximal force among Healthy groups at either timepoint. While force was diminished amongst the injury groups compared to healthy TA muscles 8 dpi, CoO-TRIM treated TA forces were greater by 35% and 49% *vs.* Untreated and Saline mice, respectively (**Figure 1A**). At 14 dpi, significant losses remained in forces within Untreated and Saline groups, whereas injured TA muscles injected with CoO-TRIM recovered to healthy TA forces, including a 22% greater force *vs.* Untreated and Saline injured muscles (**Figure 1B**). There were no significant differences at 8- or 14 dpi between Untreated and Saline mice following injury.

**Figure 1.**
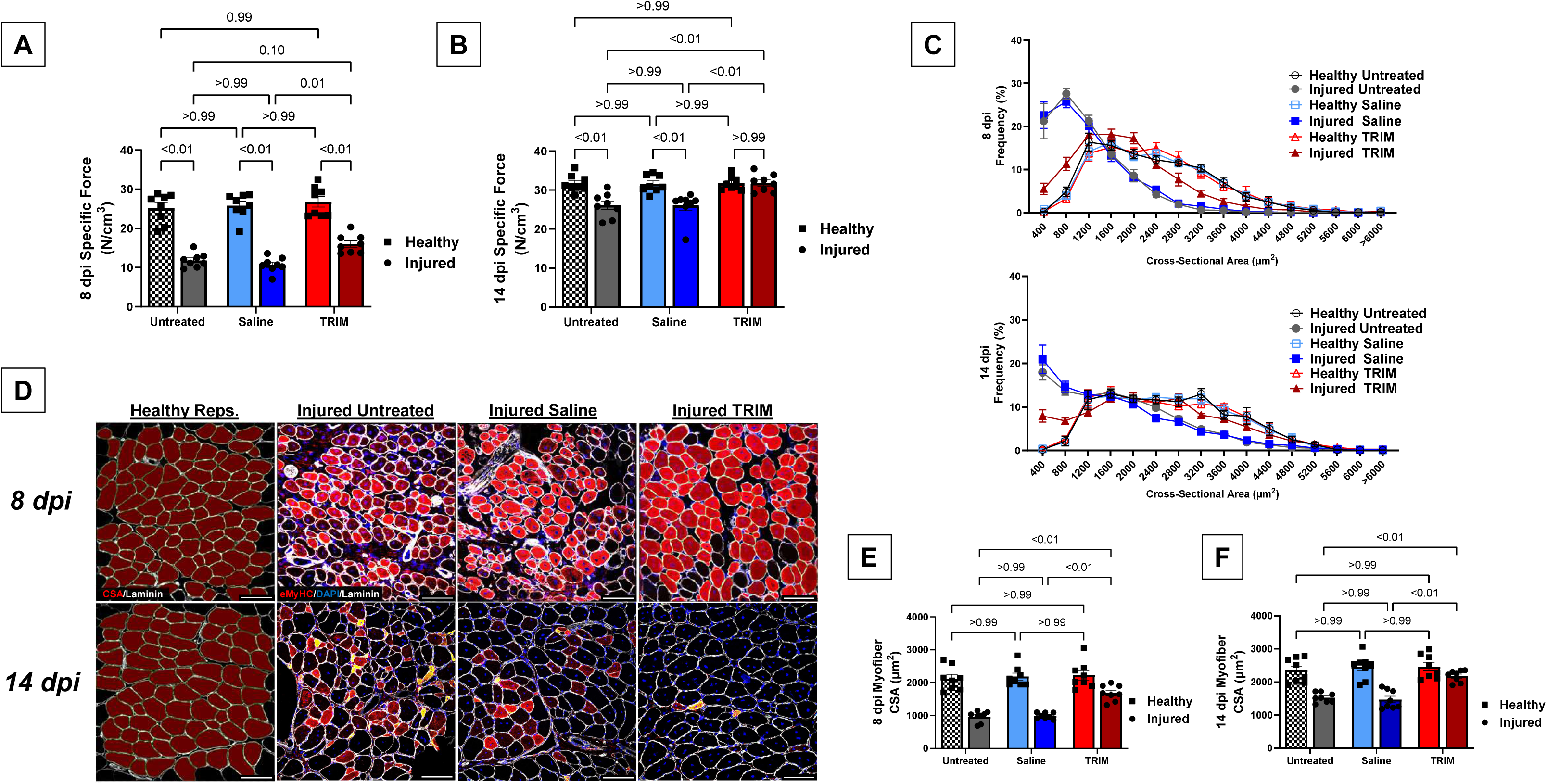
CoO-TRIM treatment increased functional recovery and muscle fiber size following injury. Summary values for max specific force via direct muscle stimulation at 120 hz at **A**) 8 dpi and **B**) 14 dpi. **C**) Muscle fiber CSA frequency distribution 8- and 14 dpi. **D**) Representative images of TA muscle cross-sections, *Left column;* representative images for all healthy controls, *Top* 8 dpi and *Bottom* 14 dpi. Laminin (white) identified basal laminae surrounding fibers, which were analyzed for CSA (red). Injured representative images stained for; Laminin (white), eMyHC (red) identified actively regenerated fibers analyzed for CSA, DAPI (blue) nuclei. For injured TAs, only eMyHC+ and CLN+ fibers were analyzed compared to healthy groups. Summary values of mean fiber CSA, **E**) 8 dpi and **F**) 14 dpi. Summary values are means ± SEM (n=8/group). Comparisons made by 2-Way ANOVA (condition x treatment), p<0.05 = significant. Scale bars = 100 µm

### Muscle Fiber CSA is Enhanced with CoO-TRIM Treatment

Fiber CSA was partitioned into 400 µm^2^ increments to create relative frequency histogram plots (**Figure 1C**). No changes were observed in fiber CSA or frequency distribution between healthy control groups 8- or 14 dpi (**Figure 1C, 1E, 1F**). Following CoO-TRIM treatment in injured TAs 8 dpi, fiber CSA was greater by 74% *vs*. Untreated TAs and 70% *vs.* Saline treated TAs (**Figure 1E**). The greater fiber CSA was evident in the frequency distribution showing a rightward shift in fiber size *vs.* Untreated and Saline treated TAs (**Figure 1C**), with a reduction in the number of small area fibers (<800 µm^2^, **Figure S1A**) and an increase in large area fibers (>2000 µm^2^, **Figure S1B**) comparable to the healthy contralateral TA groups. At 14 dpi, these trends continued as fiber CSA returned to healthy TA muscle levels and was 43% greater *vs.* Untreated TAs (**Figure 1F**) and 33% greater *vs.* Saline treated TAs (**Figure 1F**).

### Muscle Regeneration is Accelerated in CoO-TRIM Treated TA Muscle

8 dpi, we observed 29% more eMyHC^+^ muscle fibers (actively regenerating muscle fibers) in injured CoO-TRIM treated TA muscles compared to Untreated TA muscles, and 30% more compared to Saline TA muscles (**Figure 2B**). Accounting for total fiber regeneration (eMyHC^+^ + central nucleated fibers (CLN) /mm^2^), CoO-TRIM TAs had 25% more regenerated muscle fibers than Untreated TAs and 24% more than Saline Treated TAs (**Figure 2C**). No changes were observed between healthy control groups or between the Untreated *vs.* Saline treated injury groups. 14 dpi, CoO-TRIM treated TA’s had a substantially lower number of eMyHC^+^ muscle fibers than the Untreated and Saline mice, with no difference in the total number of regenerated fibers indicating that CoO-TRIM accelerated muscle regeneration following injury (**Figure 2D**).

**Figure 2.**
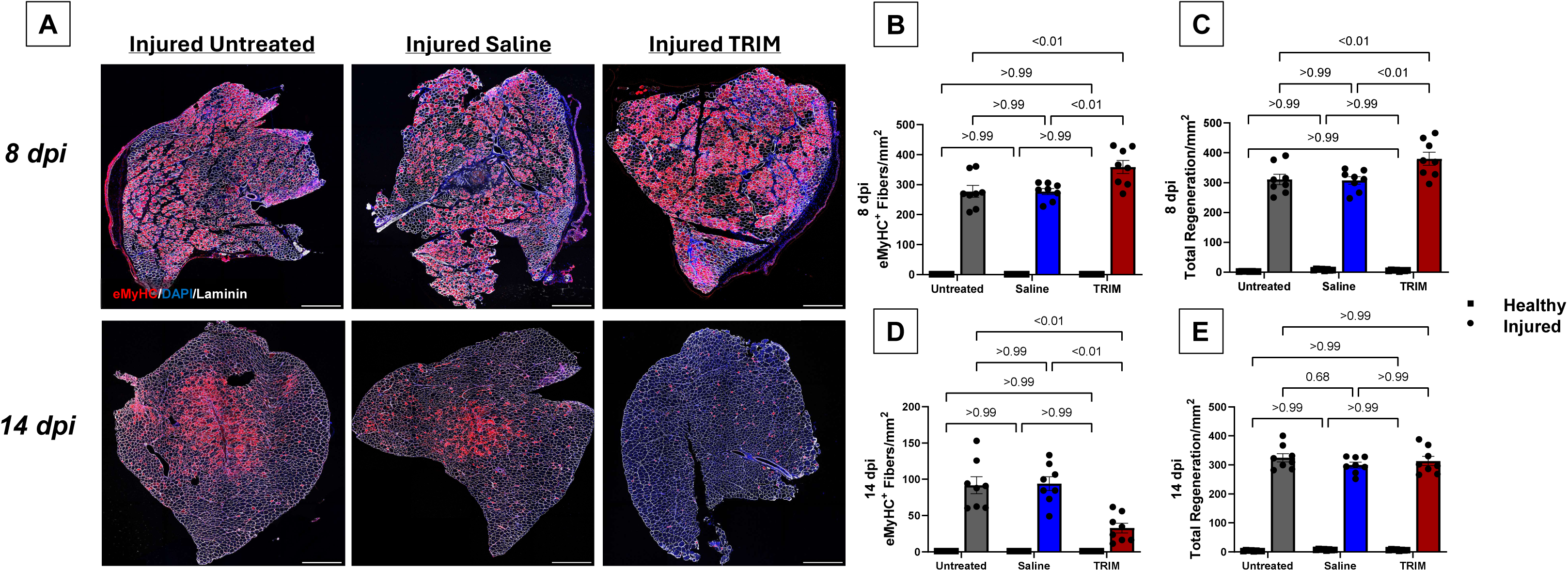
Muscle regeneration is accelerated in CoO-TRIM treated muscles following injury. **A**) Representative images of TA muscle cross-sections at *Top row,* 8 dpi and *Bottom row,* 14 dpi. Laminin (white) identified basal laminae surrounding fibers, eMyHC (red) identified actively regenerating muscle fibers, DAPI (blue) identified nuclei. Summary values comparing **B**) the number of eMyHC+ fibers and **C**) total number of regenerating muscle fibers (eMyHC+/CLN+ and eMyHC-/CLN+), normalized to muscle cross-section area (mm^2^) at 8 dpi. Summary values comparing **D**) the number of eMyHC+ fibers and **E**) total number of regenerating muscle fibers normalized to muscle cross-section area (mm^2^) at 14 dpi. Summary values are means ± SEM (n=8/group). Comparisons made by 2-Way ANOVA (condition x treatment), p<0.05 = significant. Scale bars = 500 µm

Due to the similar outcomes of the muscle function and regeneration experiments for the Untreated and Saline treated groups, only Saline and CoO-TRIM treated muscles were immunoblotted. Concomitant with immunofluorescence images, immunoblot analysis shows at 8 dpi, CoO-TRIM treated TAs had a 37% greater abundance of Pax7 (**Figure 3A Top row**), 50% more MyoD (**Figure 3B Top row**), and 47% more Myogenin (P= 0.02, **Figure 3C Top row**) than the Saline treated TAs. As indicated by the resolution of eMyHC^+^ muscle fibers in CoO-TRIM treated TAs at 14 dpi, lower amounts of Pax7 (43%, **Figure 3A Bottom row**), MyoD (56%, **Figure 3B Bottom row**), and MyoG (45%, **Figure 3C Bottom row**) were observed compared to the Saline treated TAs as myogenic factors return to near basal levels with muscle fibers completing the regenerative process.

**Figure 3.**
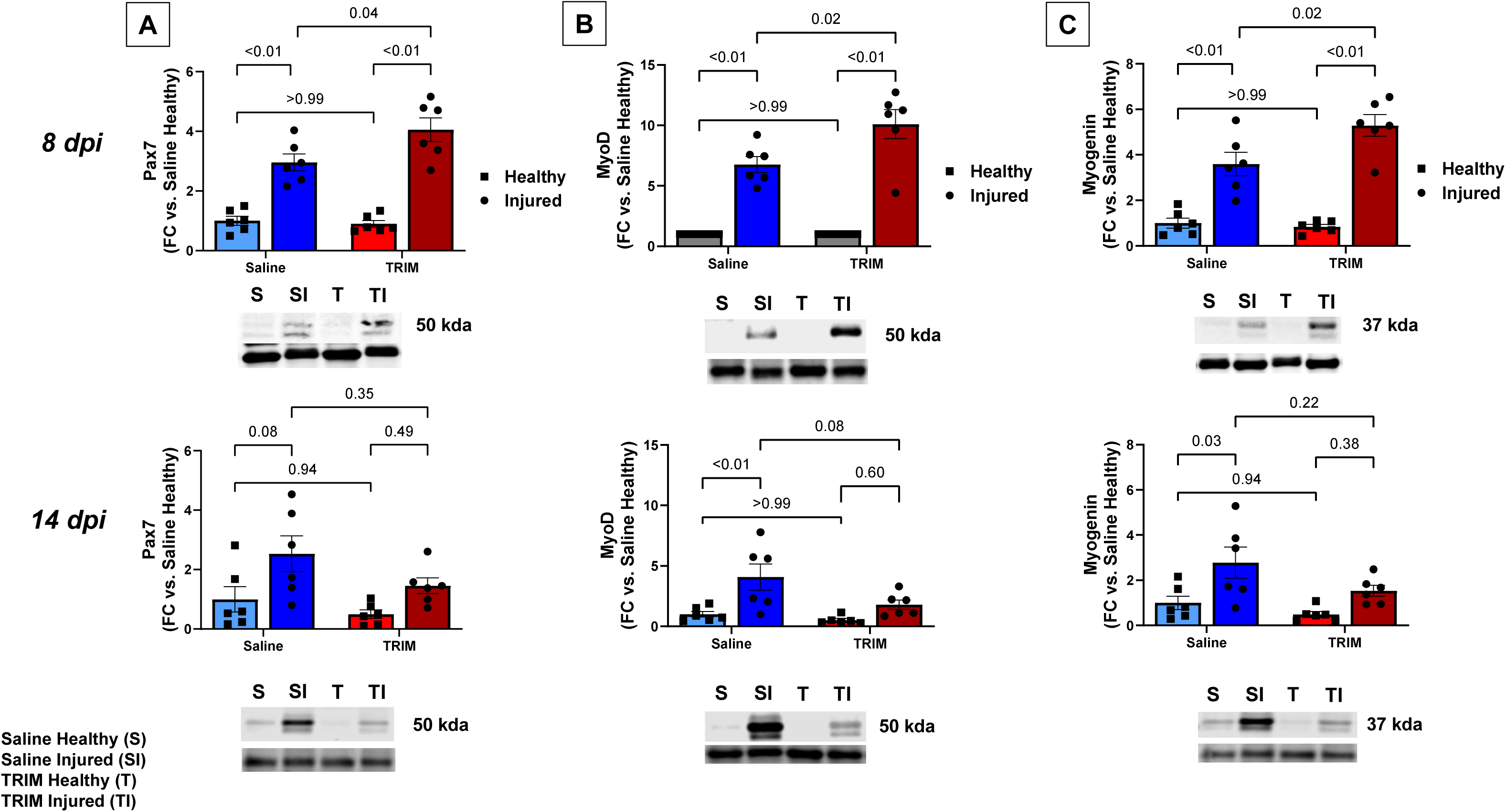
Myogenic transcription factors resolve with CoO-TRIM treatment 14 dpi. Representative immunoblots and mean densitometric data for **A**) Pax7, **B**) MyoD, and **C**) Myogenin in Saline and CoO-TRIM TA muscles *Top row,* 8 dpi and *Bottom row,* 14 dpi. (n=6/group). Comparisons made by 2-Way ANOVA (condition x treatment), p<0.05 = significant

### Muscle Anabolic Pathways Following CoO-TRIM Treatment

Given the increases in muscle force and fiber CSA in response to CoO-TRIM, we immunoblotted for the phosphorylated, total and phosphorylated to total ratio of Akt, mTOR, p70s6k and 4E-BP1 to investigate anabolic signaling contributions to structural and functional enhancements. No statistical differences were observed in pathway activation between healthy groups at 8 dpi (**Figure S3.1 A-D**) or 14 dpi (**Figure S3.2 A-D**). Injured TA muscles receiving CoO-TRIM injections had increased phosphorylated to total Akt compared to Saline treated mice and healthy contralateral controls 8 dpi (**Figure S3.1 A**), with increased phosphorylated Akt (**Figure S3.1 A**) and decreased total Akt in injured Saline and CoO-TRIM TAs (**Figure S3.1 A**). 14 dpi, phosphorylated and total Akt was elevated in injured muscles treated with Saline, with no changes observed in Akt phosphorylated to total ratio (**Figure S3.2 A**). 8 dpi, phosphorylated, total, and mTOR phosphorylated to total ratio increased in injured Saline and CoO-TRIM treated TA muscles compared to healthy contralateral controls, with no difference between injured groups (**Figure S3.1 B**). 14 dpi, mTOR signaling returned to healthy levels with no differences between healthy and injured groups, regardless of treatment (**Figure S3.2 B**). Total 4E-BP1 was increased in injured groups compared to healthy controls, with no differences in phosphorylation or the phosphorylated to total ratio 8 dpi (**Figure S3.1 C**). 14 dpi, Saline and CoO-TRIM treated injured TA muscles had increased total and phosphorylated 4E-BP1 compared to healthy controls, with increased phosphorylation to total ratio in injured Saline TA muscles (**Figure S3.2 C**). Phosphorylated and total p70s6k were increased following CoO-TRIM injection compared to healthy contralateral TA muscles 8 dpi, with no difference compared to injured Saline mice (**Figure S3.1 D**). No difference was observed between groups in phosphorylated to total p70s6k ratio (**Figure S3.1 D**). 14 dpi, phosphorylated, total, and the ratio of p70s6k returned to healthy levels in injured TA muscles injected with CoO-TRIM, while Saline treated injured muscles had elevated phosphorylation and phosphorylated to total ratio compared to injured CoO-TRIM muscles and healthy contralateral controls (**Figure S3.2 D**).

Active µ- and m-calpains (1 and 2) cleaves the skeletal muscle structural protein αII-spectrin from 240 to a 145 kDa product, while active caspase 3 produces a cleavage byproduct at 120 kDa. Therefore, we measured µ- and m-calpain and caspase 3 activity by immunoblotting for αII-spectrin 145 kDa and 120 kDa cleaved protein byproduct. While both proteolytic markers were increased in injury groups at 8- and 14 dpi, no differences were observed between Saline and TRIM groups (**Figure S4 A-D**).

### CoO-TRIM Treatment Promotes Angiogenesis

The ion release following matrix degradation has been observed to promote angiogenic properties enhancing the regeneration of soft tissue. We evaluated CD31 in muscle cross sections as a biomarker of endothelial cells and vessel growth. As expected, post-injury, Untreated, Saline and CoO-TRIM injected TAs experienced increased angiogenesis *vs.* Contralateral controls at the 8- and 14-day timepoints with revascularization taking place. While all injury groups exhibit increased intramuscular vascularity, CoO-TRIM injected TAs had a greater angiogenic response up to 14 dpi *vs.* Untreated and Saline treated TAs (**Figure 4C and 4D**). No changes were observed in vessel area between healthy contralateral limbs.

**Figure 4.**
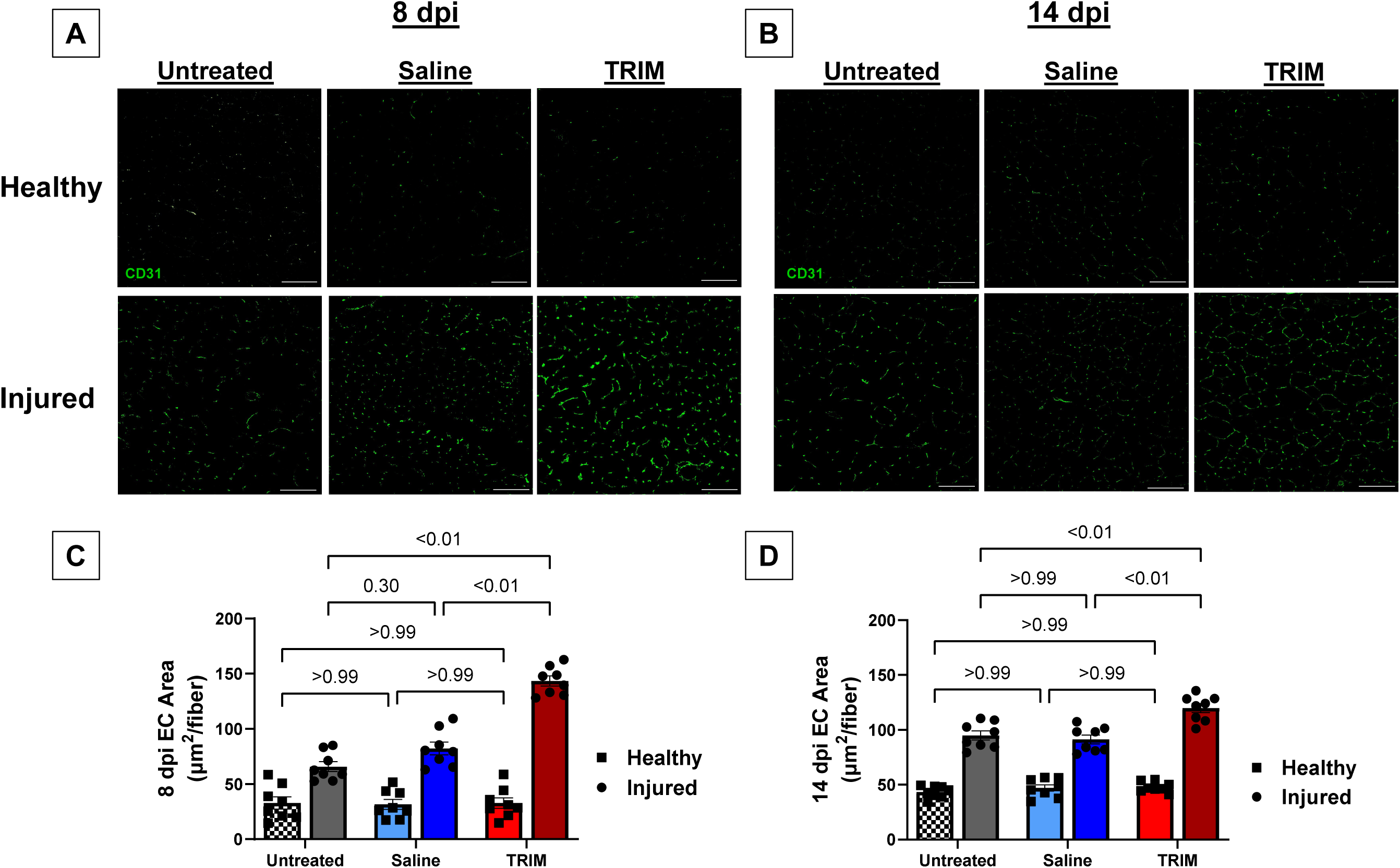
CoO-TRIM enhances muscle revascularization. Representative images of TA muscle cross-sections at **A**) 8 dpi and **B**) 14 dpi. CD31+ immunostaining (green) identifies endothelial cells (EC) as a measure of vascularity. EC area was calculated as: total EC area/ # of fibers. Summary values for **C**) 8 dpi and **D**) 14 dpi are means ± SEM (n=8/group). Comparisons made by 2-Way ANOVA (condition x treatment), p<0.05 = significant. Scale bars = 100 µm

### Angiogenic Factor Expression Following CoO-TRIM Treatment

VEGF has been reported to be a potent regulator of regeneration, with synergistic effects on satellite cell proliferation and angiogenesis. At 8 dpi, injured TAs treated with CoO-TRIM experienced a near 2-fold increase in VEGF concentration *vs.* Untreated (P= <0.01) and Saline treated (P= <0.01) TAs, with no observed effects in healthy muscle tissue (**Figure 5B Top row**). With increased time for matrix degradation and ion release, VEGF concentration further elevated 14 dpi in CoO-TRIM treated TAs following injury (**Figure 5B Bottom row**). While significant increases were found within groups (Healthy *vs.* Injured) for *Hif1α* and IGF-1, no differences were observed between groups at 8 dpi (**Figure 5A and 5C Top row**) or 14 dpi (**Figure 5A and 5C Bottom row**).

**Figure 5.**
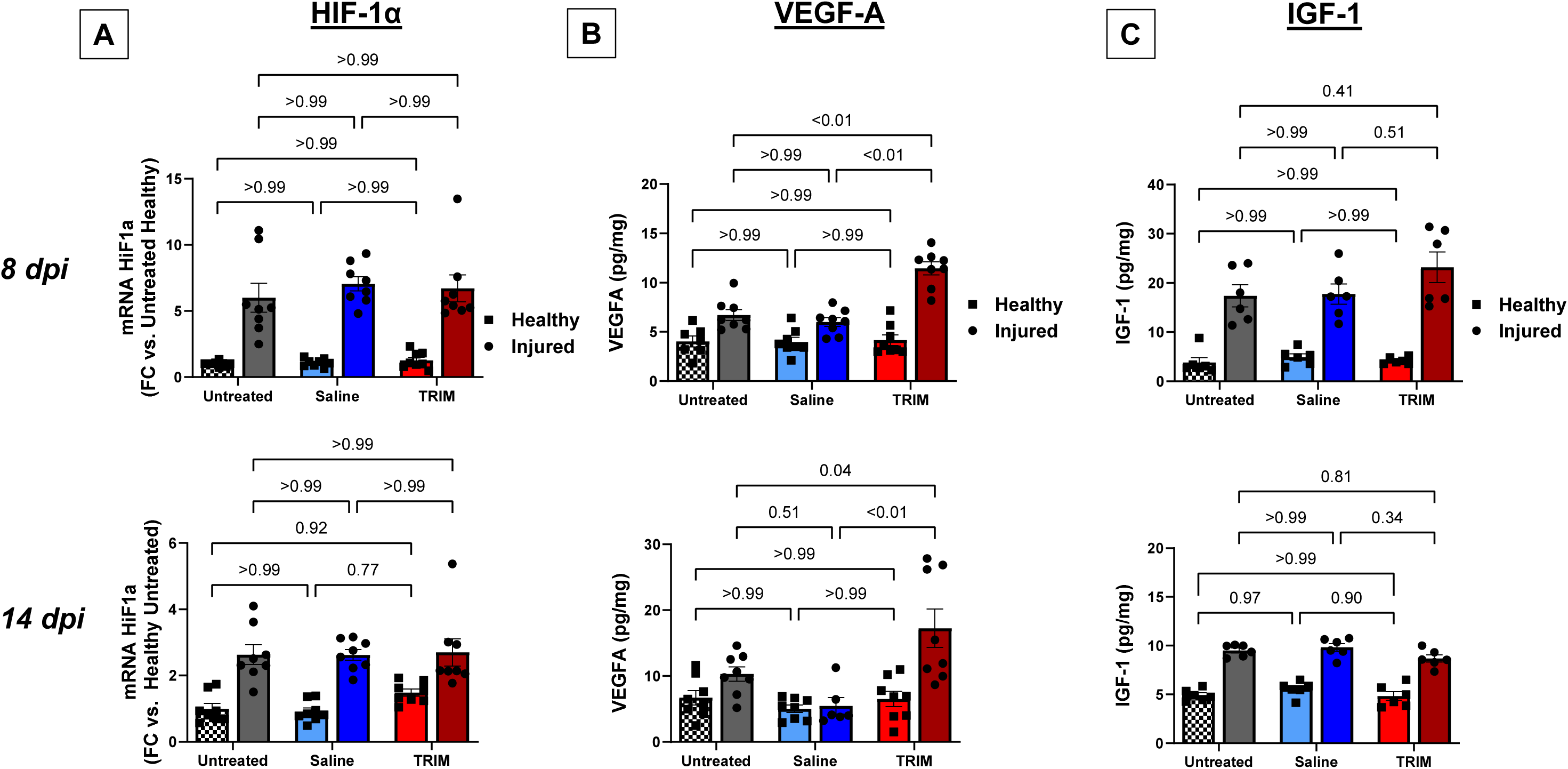
VEGF is increased with CoO-TRIM following injury. *Top row,* 8 dpi and *Bottom row,* 14 dpi. **A**) *Hif1α* gene expression in isolated TA muscle RNA assessed by RT-PCR. **B**) VEGFA and **C**) IGF-1 concentrations in whole muscle TA homogenates probed by ELISA. Summary values are means ± SEM (n=6-8/group). Comparisons made by 2-Way ANOVA (condition x treatment), p<0.05 = significant

### CoO-TRIM Increases Acute Inflammatory Response

Given the importance of early inflammatory cell infiltration during the early phase of regeneration, we immunoblotted for immune cell marker CD45. 8 dpi following CoO-TRIM treatment, CD45 was elevated in injured TAs *vs* Saline Treated (**Figure S6A**), with no significant difference observed at 14 dpi (**Figure S6B**). No inflammatory responses were observed following CoO-TRIM treatment in healthy WT muscle (**Figure 6 and S7**). At 8 dpi, CXCL1 (**Figure 6A Top**), IL-6 (**Figure 6B Top**), TNF-α (**Figure 6C Top**), and IL-10 (**Figure 6E Top**) were elevated in CoO-TRIM injected TA muscles *vs.* Untreated and Saline injected TAs post-injury. IL-1β was reduced in Saline and CoO-TRIM TA muscles *vs.* Untreated 8 dpi (**Figure 6D Top)**. At 14 dpi, no differences were observed in CXCL1, IL-6, TNF-α, or IL-1β between injured groups (**Figure 6A-E Bottom)**. IL-10, a regenerative cytokine released by M2 macrophages, remained elevated in CoO-TRIM injected TAs at 14 dpi (**Figure 6E Bottom**). There were no differences in the remaining cytokines analyzed (**Figure S7 A-E**).

**Figure 6.**
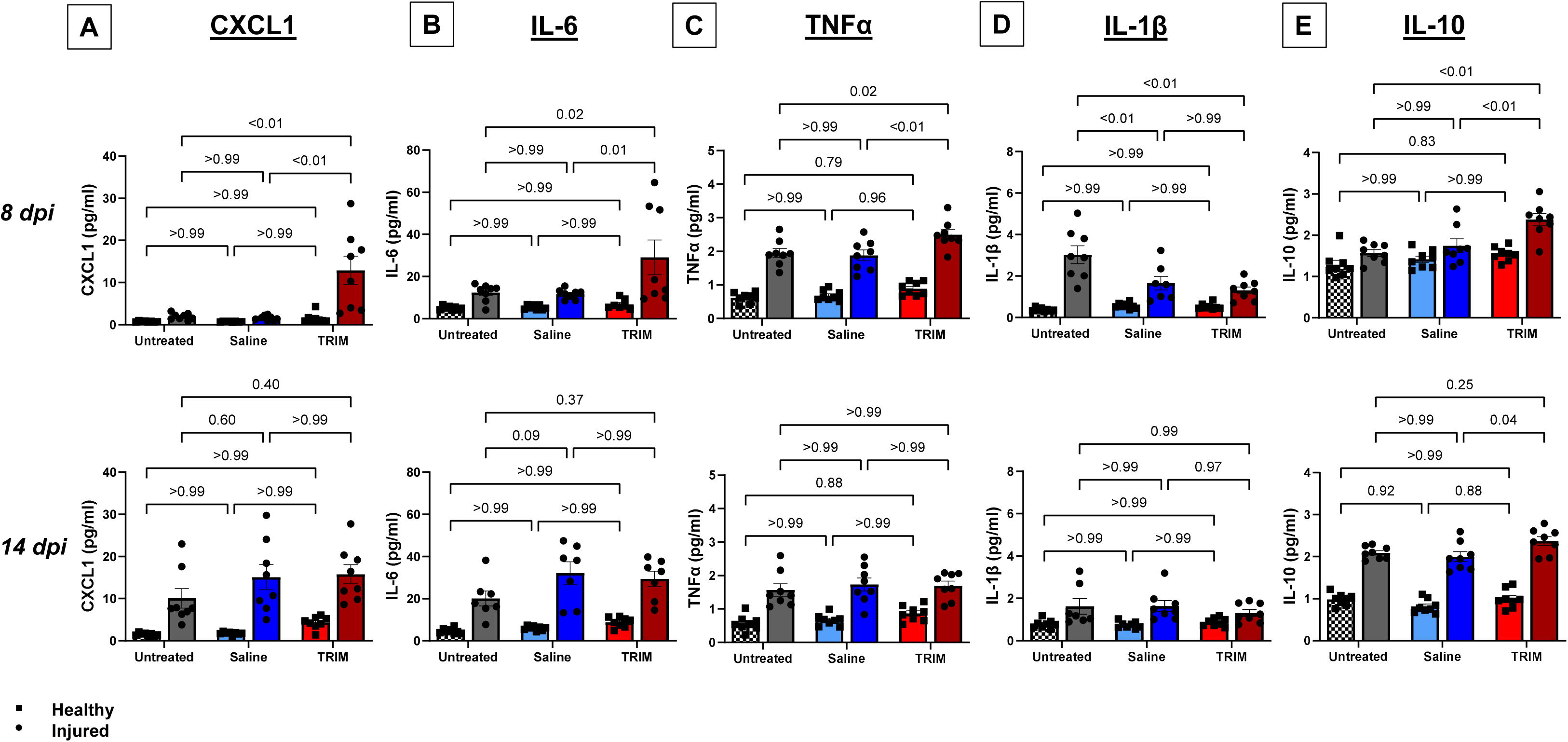
CoO-TRIM stimulates early inflammatory response following injury. Chemokine and cytokine concentration *Top,* 8 dpi and *Bottom,* 14 dpi for **A**) CXCL1, **B**) IL-6, **C**) TNF-α, **D**) IL-1β, and **E**) IL-10. Summary values are means ± SEM (n=7-8/group). Comparisons made by 2-Way ANOVA (condition x treatment), p<0.05 = significant

## Discussion

Injectable borophosphate ion matrices are promising inorganic biomaterials, capable of accelerating the regeneration of damaged muscle structure, function, and angiogenesis in accord with elevated markers of local inflammatory cytokines. While a previous group investigated an aluminoborate inorganic biomaterial in a model of volumetric muscle loss (VML), the present study is the first to treat an acute muscle injury with a cobalt-incorporated borophosphate, reporting upon prominent phases of the regeneration cycle. Herein we report when BaCl_2_-injured C57BL/6J mice were injected with CoO-TRIM: 1) Max isometric force was recovered by 14 dpi, 2) muscle fiber regeneration and myogenic factors were accelerated, 3) regenerating muscle fibers were larger, 4) intramuscular vascularity was increased, 5) VEGF protein concentration was enhanced, and 6) CXCL1, IL-6, TNF-α, and IL-10 cytokine concentrations were increased 8 dpi, with IL-10 remaining elevated 14 dpi. No changes were observed in Healthy contralateral controls at either timepoint.

### Skeletal Muscle Regeneration with CoO-TRIM Treatment

Mild injury to skeletal muscle causes ultrastructural damage, reducing contractile force, prompting a series of overlapping degenerative and regenerative events, culminating in myofiber regeneration(31, 32). Local CoO-TRIM treatment following acute muscle injury accelerated the myogenic cycle, linking satellite cell activity to muscle fiber maturation and functional restoration. At 8 dpi, CoO-TRIM injected muscles exhibited increased satellite cell abundance with concomitant MyoD and myogenin expression, indicative of active proliferation and differentiation. Markers of mTOR activation were elevated in CoO-TRIM injected muscles 8 dpi relative to Healthy muscles, but not significantly different compared to Saline injected muscles suggesting CoO-TRIM does not negatively regulate hypertrophic signaling necessary for myofiber repair. By 14 dpi, actively regenerating myofibers had decreased threefold following CoO-TRIM injection, coinciding with complete restoration of contractile force compared to a 22% deficit observed in Untreated and Saline injected TA muscles. Early maturation is functionally significant as delayed fiber recovery correlates with prolonged muscle weakness and reduced tissue resilience post-injury. Satellite cell expansion and differentiation factors (MyoD and myogenin) were no longer different from uninjured muscle following CoO-TRIM treatment by 14 dpi while remaining elevated in Saline treated groups, suggesting significant acceleration of regeneration without overactivation. In contrast, CoO-TRIM did not exert any detectable effects on muscle function, regeneration, or fiber size in uninjured muscle, indicating a need for activation of inflammatory pathways. Investigations conducted by our group treating dystrophic mouse muscle demonstrated enhanced muscle function and fiber regeneration following CoO-TRIM treatment, but no changes in WT, healthy muscle, while a similar aluminoborate TRIM in VML enhanced muscle regeneration within the tissue void compared to controls(30, 33). Together, these investigations suggest inorganic borophosphate TRIMs may be effective treatments enhancing the native myogenic response without the addition of exogenous cells or growth factors.

### Increased VEGF and Angiogenesis after CoO-TRIM Injection

Revascularization following muscle injury precedes myogenesis with the restoration of the blood supply necessary for the growth of mature fibers, supporting a regenerative microenvironment(34). While angiogenesis has been identified as an important therapeutic target to improve regeneration, significant challenges have prevented a therapeutic agent from successfully transitioning to the clinic(20, 35, 36). The ionic dissolution products from inorganic TRIM’s consistently demonstrate stimulation of VEGF and subsequent angiogenesis *in vitro* and *in vivo* in VML and disease(29, 30, 33). Up to 14 dpi, CoO-TRIM enhanced angiogenesis with sustained VEGF expression, promoting capillary formation and vascular remodeling. *Hif1α* levels were elevated but not significantly different from Untreated and Saline injected muscle, suggesting VEGF-driven angiogenesis was largely regulated by immune and paracrine signaling. Angiogenesis is closely coupled with myogenesis as differentiating satellite cells release pro-angiogenic growth factors, stabilizing sprouting endothelial cells. In concert, vascular expansion supports heightened metabolic demands of proliferating myoblasts. Elevated VEGF expression following CoO-TRIM injection ensures adequate perfusion to the local muscle microenvironment even as regeneration resolves. Moreover, coordinated increases in angiogenesis and myogenesis observed in CoO-TRIM injected injured muscles coincides with reports of VEGF elevation and angiogenesis accelerating functional muscle repair.

### Modulated Inflammatory Response Following CoO-TRIM Treatment

Early regenerative events in skeletal muscle require activation of resident immune cells triggering chemotactic responses of the adaptive immune system(5, 6). Early immune cells, namely intramuscular macrophages, release proinflammatory cytokines which support clearance of muscle debris. As regeneration progresses, signaling transitions towards a pro-regenerative response, supporting efficient muscle repair(37, 38). Cytokine secretion, in particular TNF-α, IL-6, and IL-10, trigger the release of growth factors, VEGF and IGF-1 during muscle repair, increasing satellite cell proliferation and vascular remodeling(7, 8, 39). In contrast, reduced inflammation impairs muscle recovery, indicating the necessity of a local immune response at the site of injury(40). The present experiments are the first to investigate inflammatory responses following borophosphate TRIM treatment in injured muscle. We observed that CoO-TRIM modulated the inflammatory response following injury, with no observable activity in healthy tissue. In response to CoO-TRIM treatment, early elevations in CXCL1 and pro-inflammatory cytokines (IL-6, and TNF-α) were observed, consistent with enhanced immune cell recruitment and clearance of necrotic debris. By 14 dpi, these cytokines had resolved their acute elevation to similar levels measured in Untreated and Saline injected muscles, yet IL-10, a cytokine associated with M2 macrophage polarization and tissue regeneration remained elevated following CoO-TRIM treatment. This dynamic immune response suggests that CoO-TRIM may support a tightly regulated transition from degenerative inflammatory signaling to regenerative signaling. Notably, the persistence of IL-10 may provide a favorable milieu for sustained angiogenesis and satellite cell differentiation, consistent with the increases in VEGF expression, vessel density, and myogenic maturation observed in CoO-TRIM injected muscles. Furthermore, immune cells contribute to metabolic support within regenerating tissue, as M2 macrophages provide growth factors and cytokines that not only regulate cellular proliferation, but modulate metabolism in satellite cells and endothelial cells matching energy demands required during rapid tissue expansion. Our previous investigations in dystrophic skeletal muscle are similar, CoO-TRIM increased the already prevalent immune signaling seen in injured and diseased muscle(33). However, in healthy muscle injected with CoO-TRIM with no previous immune modulation, no changes were observed with CoO-TRIM, suggesting CoO-TRIM likely acts through preexisting inflammatory responses for downstream effects on angiogenesis and muscle fiber regeneration. Together, these findings highlight the potential for inorganic biomaterials to not only support structural regeneration but the orchestration of cellular and molecular events underlying functional recovery of injured skeletal muscle.

## Conclusion

The present work expands the application of inorganic biomaterials in regenerative medicine, demonstrating how CoO-TRIM promotes skeletal muscle regeneration through a coordinated triad of myogenesis, angiogenesis, and inflammatory signaling. This distinguishes CoO-TRIM from current strategies relying on cell-based therapies and engineered growth factor delivery. Rather, the controlled release of bioactive ions from CoO-TRIM integrates biophysical and biochemical cues to accelerate repair while minimizing off-target effects. Satellite cell activity and anabolic signaling peak at 8 days, followed by resolution of active regeneration, fiber maturation, and functional recovery by 14 days. Angiogenesis persists to maintain vascular support, while immune modulation prompts inflammatory cues to prime and sustain the regenerative microenvironment without excessive inflammation. These findings highlight CoO-TRIM as an alternative therapy for skeletal muscle repair, offering a scalable, cell and growth factor free approach to enhancing muscle recovery following injury.

## Supporting information

Western Blot Supplement

## Author Contributions

Conceptualization: J.A.K., A.B.M., Data Curation: J.A.K., A.B.M., Formal Analysis: J.A.K., A.B.M., Funding Acquisition: J.A.K., A.B.M., R.K.B., R.L.B., Investigation: J.A.K., A.B.M., Methodology: J.A.K., A.B.M., Project Administration: J.A.K., A.B.M., Data Collection: J.A.K., J.H., D.R.H., A.G.N., S.D.G., L.D.G., Resources: A.B.M., R.K.B., Validation: J.A.K., A.B.M., Visualization: J.A.K., A.B.M., Writing – original draft: J.A.K., Writing – review and editing: J.A.K., J.H., D.R.H., R.L.B., A.G.N., S.D.G., L.D.G., R.K.B., A.B.M.. All authors have read and approved the final version of this manuscript and agree to be accountable for all aspects of the work ensuring that questions related to accuracy or integrity of any part of the work are appropriately investigated and resolved. All persons designated as authors qualify for authorship and all those who qualify for authorship are listed.

## Data Availability

The data that support the findings of this study are available from the corresponding author upon reasonable request.

## Acknowledgments

The authors would like to acknowledge the following organizations for providing funding support in completion of this work; Sydney and J.L. Huffines Institute for Sports Medicine and Human Performance (J.A.K.), Texas A&M School of Education and Human Development (J.A.K., A.B.M.), Texas A&M Department of Kinesiology and Sports Management (J.A.K., A.B.M.), the Ignition Grant Initiative at Missouri S&T (R.K.B., R.L.B.), and a National Institutes of Health Loan Repayment Award through NIAMS (A.B.M.).

## Ethical Publication Statement

We confirm that we have read the Journal’s position on issues involved in ethical publication and affirm that this report is consistent with those guidelines. All protocols in this study were approved by Animal Care and Use Committees at Texas A&M University (AUP 2022-0215). Animal Care was in accordance with the National Research Council’s *Guide for the Care and Use of Laboratory Animals (Eighth Edition, 2011)*.

## References

1. Molloy JM, Pendergrass TL, Lee IE, Chervak MC, Hauret KG, Rhon DI. Musculoskeletal Injuries and United States Army Readiness Part I: Overview of Injuries and their Strategic Impact. Military Medicine. 2020;185(9-10):e1461–e71.

2. Goes RA, Lopes LR, Cossich VRA, de Miranda VAR, Coelho ON, do Carmo Bastos R, et al. Musculoskeletal injuries in athletes from five modalities: a cross-sectional study. BMC Musculoskeletal Disorders. 2020;21(1):122.

3. Teyhen DS, Goffar SL, Shaffer SW, Kiesel K, Butler RJ, Tedaldi A-M, et al. Incidence of Musculoskeletal Injury in US Army Unit Types: A Prospective Cohort Study. Journal of Orthopaedic & Sports Physical Therapy. 2018;48(10):749–57.

4. Järvinen TA, Järvinen M, Kalimo H. Regeneration of injured skeletal muscle after the injury. Muscles Ligaments Tendons J. 2013;3(4):337–45.

5. Yang W, Hu P. Skeletal muscle regeneration is modulated by inflammation. J Orthop Translat. 2018;13:25–32.

6. Tidball JG. Regulation of muscle growth and regeneration by the immune system. Nat Rev Immunol. 2017;17(3):165–78.

7. Lu CY, Santosa KB, Jablonka-Shariff A, Vannucci B, Fuchs A, Turnbull I, et al. Macrophage-Derived Vascular Endothelial Growth Factor-A Is Integral to Neuromuscular Junction Reinnervation after Nerve Injury. J Neurosci. 2020;40(50):9602–16.

8. Fantin A, Vieira JM, Gestri G, Denti L, Schwarz Q, Prykhozhij S, et al. Tissue macrophages act as cellular chaperones for vascular anastomosis downstream of VEGF-mediated endothelial tip cell induction. Blood. 2010;116(5):829–40.

9. Liu S, Yu JM, Gan YC, Qiu XZ, Gao ZC, Wang H, et al. Biomimetic natural biomaterials for tissue engineering and regenerative medicine: new biosynthesis methods, recent advances, and emerging applications. Mil Med Res. 2023;10(1):16.

10. Lei B, Guo B, Rambhia KJ, Ma PX. Hybrid polymer biomaterials for bone tissue regeneration. Front Med. 2019;13(2):189–201.

11. Rossi CA, Flaibani M, Blaauw B, Pozzobon M, Figallo E, Reggiani C, et al. In vivo tissue engineering of functional skeletal muscle by freshly isolated satellite cells embedded in a photopolymerizable hydrogel. Faseb j. 2011;25(7):2296–304.

12. Kim JH, Ko IK, Atala A, Yoo JJ. Progressive Muscle Cell Delivery as a Solution for Volumetric Muscle Defect Repair. Sci Rep. 2016;6:38754.

13. Goldman SM, Henderson BEP, Walters TJ, Corona BT. Co-delivery of a laminin-111 supplemented hyaluronic acid based hydrogel with minced muscle graft in the treatment of volumetric muscle loss injury. PLoS One. 2018;13(1):e0191245.

14. Beauchamp JR, Morgan JE, Pagel CN, Partridge TA. Dynamics of Myoblast Transplantation Reveal a Discrete Minority of Precursors with Stem Cell–like Properties as the Myogenic Source. Journal of Cell Biology. 1999;144(6):1113–22.

15. Boonen KJ, Post MJ. The muscle stem cell niche: regulation of satellite cells during regeneration. Tissue Eng Part B Rev. 2008;14(4):419–31.

16. Montarras D, Morgan J, Collins C, Relaix F, Zaffran S, Cumano A, et al. Direct isolation of satellite cells for skeletal muscle regeneration. Science. 2005;309(5743):2064–7.

17. Partridge TA, Morgan JE, Coulton GR, Hoffman EP, Kunkel LM. Conversion of mdx myofibres from dystrophin-negative to -positive by injection of normal myoblasts. Nature. 1989;337(6203):176–9.

18. Raimondo TM, Li H, Kwee BJ, Kinsley S, Budina E, Anderson EM, et al. Combined delivery of VEGF and IGF-1 promotes functional innervation in mice and improves muscle transplantation in rabbits. Biomaterials. 2019;216:119246.

19. Borselli C, Storrie H, Benesch-Lee F, Shvartsman D, Cezar C, Lichtman JW, et al. Functional muscle regeneration with combined delivery of angiogenesis and myogenesis factors. Proc Natl Acad Sci U S A. 2010;107(8):3287–92.

20. Anderson EM, Silva EA, Hao Y, Martinick KD, Vermillion SA, Stafford AG, et al. VEGF and IGF Delivered from Alginate Hydrogels Promote Stable Perfusion Recovery in Ischemic Hind Limbs of Aged Mice and Young Rabbits. J Vasc Res. 2017;54(5):288–98.

21. Lee KY, Peters MC, Anderson KW, Mooney DJ. Controlled growth factor release from synthetic extracellular matrices. Nature. 2000;408(6815):998–1000.

22. Mitchell AC, Briquez PS, Hubbell JA, Cochran JR. Engineering growth factors for regenerative medicine applications. Acta Biomater. 2016;30:1–12.

23. Day RM. Bioactive glass stimulates the secretion of angiogenic growth factors and angiogenesis in vitro. Tissue Eng. 2005;11(5-6):768–77.

24. Ege D, Zheng K, Boccaccini AR. Borate Bioactive Glasses (BBG): Bone Regeneration, Wound Healing Applications, and Future Directions. ACS Appl Bio Mater. 2022;5(8):3608–22.

25. Hench LL, Paschall HA. Direct chemical bond of bioactive glass-ceramic materials to bone and muscle. J Biomed Mater Res. 1973;7(3):25–42.

26. Ciraldo FE, Boccardi E, Melli V, Westhauser F, Boccaccini AR. Tackling bioactive glass excessive in vitro bioreactivity: Preconditioning approaches for cell culture tests. Acta Biomater. 2018;75:3–10.

27. Liu X, Rahaman MN, Day DE. Conversion of melt-derived microfibrous borate (13-93B3) and silicate (45S5) bioactive glass in a simulated body fluid. J Mater Sci Mater Med. 2013;24(3):583–95.

28. Freudenberger PT, Blatt RL, Brow RK. Dissolution rates of borophosphate glasses in deionized water and in simulated body fluid. Journal of Non-Crystalline Solids: X. 2023;18:100181.

29. Bromet BA, Blackwell NP, Abokefa N, Freudenberger P, Blatt RL, Brow RK, et al. The angiogenic potential of pH-neutral borophosphate bioactive glasses. J Biomed Mater Res A. 2023;111(10):1554–64.

30. Jia W, Hu H, Li A, Deng H, Hogue CL, Mauro JC, et al. Glass-activated regeneration of volumetric muscle loss. Acta Biomater. 2020;103:306–17.

31. Lovering RM, De Deyne PG. Contractile function, sarcolemma integrity, and the loss of dystrophin after skeletal muscle eccentric contraction-induced injury. Am J Physiol Cell Physiol. 2004;286(2):C230–8.

32. Laumonier T, Menetrey J. Muscle injuries and strategies for improving their repair. J Exp Orthop. 2016;3(1):15.

33. Morton A, Kendra J, Naman A, Blatt R, Brow R, Segal S, et al. Time Release Ion Matrix Regenerates Dystrophic Skeletal Muscle. Res Sq. 2025.

34. Jacobsen NL, Morton AB, Segal SS. Angiogenesis precedes myogenesis during regeneration following biopsy injury of skeletal muscle. Skeletal Muscle. 2023;13(1):3.

35. Tillman B, Hardin-Young J, Shannon W, Russell AJ, Parenteau NL. Meeting the need for regenerative therapies: translation-focused analysis of U.S. regenerative medicine opportunities in cardiovascular and peripheral vascular medicine using detailed incidence data. Tissue Eng Part B Rev. 2013;19(2):99–115.

36. Silvestre JS, Smadja DM, Lévy BI. Postischemic revascularization: from cellular and molecular mechanisms to clinical applications. Physiol Rev. 2013;93(4):1743–802.

37. Rigamonti E, Zordan P, Sciorati C, Rovere-Querini P, Brunelli S. Macrophage plasticity in skeletal muscle repair. Biomed Res Int. 2014;2014:560629.

38. Latroche C, Weiss-Gayet M, Muller L, Gitiaux C, Leblanc P, Liot S, et al. Coupling between Myogenesis and Angiogenesis during Skeletal Muscle Regeneration Is Stimulated by Restorative Macrophages. Stem Cell Reports. 2017;9(6):2018–33.

39. Bernard C, Zavoriti A, Pucelle Q, Chazaud B, Gondin J. Role of macrophages during skeletal muscle regeneration and hypertrophy-Implications for immunomodulatory strategies. Physiol Rep. 2022;10(19):e15480.

40. Shireman PK, Contreras-Shannon V, Ochoa O, Karia BP, Michalek JE, McManus LM. MCP-1 deficiency causes altered inflammation with impaired skeletal muscle regeneration. J Leukoc Biol. 2007;81(3)y:775–85.

