## Supplement Text for "Skeletal Muscle Regeneration is Accelerated Following Injection of Time Release Ion Matrix in Injured Mice"

**Supplement Material**

**Methods**

*Biomaterial Fabrication*

The raw materials were mixed, placed in a platinum crucible calcined overnight at 300°C to evolve water and $CO_2$, then melted at 1150°C for 30 minutes, stirred with a platinum rod, then melted for another 30 minutes. The homogeneous melt was poured into graphite molds, then annealed at 350°C for 1 h before cooling to room temperature. Annealed CoO-TRIM was ground to form the CoO-TRIM particles with a diameter <10 microns using a Spex mill and placed in 1.5 mL microcentrifuge tubes in a desiccator to store prior to use.

*Muscle Force*

The distal tendon of the TA muscle was isolated, secured with a 2-0 suture, then severed from its insertion. The mouse was placed prone on a platform set to 37°C using a heated water pump; a heat lamp maintained the muscle at 32°C. A vertical metal peg was inserted into the patellar tendon to immobilize the knee, while the TA tendon was attached to a lever arm with a micrometer to adjust the optimal length ($L_o$), as determined during twitch contractions at 1 Hz. A strip of KimWipe® was wrapped around the exposed TA muscles and irrigated with sterile saline warmed to 37°C. For direct field stimulation, a pair of platinum/iridium wire electrodes were placed across the TA muscle belly to generate sufficient current to depolarize muscle fibers.

*Immunofluorescence Analysis*

TA muscles were embedded in Tissue-Plus™ O.C.T. Compound (Scigen, Fischer Scientific, Hampton, NJ, USA), frozen in liquid nitrogen-cooled 2-methyl butane (Thermo Fisher Scientific, Waltham, MA, USA) and sectioned (thickness, 10 μm) in a Cryostar NX50 Cryostat (Epredia, Kalamazoo, MI, USA) at -17°C onto a microscope slide. Sections were permeabilized in 0.5% Triton X-100, washed in Tris-buffered saline (TBS), fixed with ice-cold 4% paraformaldehyde for 10 minutes, washed 3x in TBS, then blocked with 3% bovine serum albumin (BSA)/5% normal goat serum (NGS)/3% Triton x-100 for 1 h at room temperature (RT). Primary antibodies were incubated for 60 minutes at RT in blocking buffer (as above). Sections were washed 3x in TBS, incubated with secondary antibodies in blocking buffer for 60 minutes at RT, washed 3x in TBS, and mounted in Invitrogen™ ProLong™ Gold antifade reagent with DAPI (Cat.# P36941, Fisher Scientific, Hampton, NJ, USA).

Slides were imaged on a Stellaris 5 White Light Laser confocal microscope (Leica Microsystems, Deer Park, IL, USA) using Leica LAS_X software (RRID: SCR_013673, Leica Microsystems, Deer Park, IL, USA).

*Western Blot Analysis*

All primary and secondary antibodies use a 1:1000 and 1:20,000 concentration, respectively, unless otherwise noted. Membranes were scanned with a Li-Cor Odyssey DLx Imager (Li-Cor Biotechnology, Lincoln, NE, US) and analyzed using Image Studio Lite. Primary antibodies of interest were; Pax7 (RRID: AB_528428, Cat. # PAX7, Developmental Studies Hybridoma Bank, The University of Iowa Department of Biology; Iowa City, IA, USA; Primary 1:500), MyoD (G-1) (RRID: AB_2813894, Cat. # sc-377460, Santa Cruz Biotechnology, Santa Cruz, CA; Primary 1:500), Myogenin (F5D) (RRID: AB_627980, Cat. # sc-377460, Santa Cruz Biotechnology, Santa Cruz, CA; Primary 1:500), phospho-Akt (Ser473) (RRID: AB_329825, Cat.# 9271; Cell Signaling Technology, Danvers, MA), Akt1 (RRID: AB_915788, Cat.# 2938; Cell Signaling Technology, Danvers, MA), phospho-mTOR (Ser2448) (RRID: AB_10691552, Cat.# 5536; Cell Signaling Technology, Danvers, MA), mTOR (RRID: AB_2105622, Cat.# 2983; Cell Signaling Technology, Danvers, MA), phosphor-p70S6K (THr389) (RRID: AB_330944, Cat.# 9205; Cell Signaling Technology, Danvers, MA), p70S6K (RRID: AB_331676, Cat.# 9202; Cell Signaling Technology, Danvers, MA), phospho-4E-BP1 (Thr37/46) (RRID: AB_560835, Cat.# 2855; Cell Signaling Technology, Danvers, MA), 4eBP1 (RRID: AB_2097841, Cat.# 9644; Cell Signaling Technology, Danvers, MA), Alpha II spectrin (RRID: AB_2194351, Cat.# sc-48382; Santa Cruz Biotechnology, Santa Cruz, CA, Primary 1:250 incubated overnight at 4 °C then 3 h at RT, Secondary 1:5,000), CD45 (RRID: AB_442810, Cat. # ab10558, Abcam, Waltham, MA, USA, Primary 1:500, Secondary 1:5,000).

*ELISA*

Concentrations for VEGFA, IGF-1, and chemokine/cytokines were determined by comparing samples to respective recombinant protein standards supplied by the manufacturer. For VEGFA and IGF-1, analyte concentrations were normalized to individual sample protein concentration assessed using the Bradford method (Sigma-Aldrich, St. Louis, MO, USA) (pg/mg). For Proinflammatory panel chemokine/cytokine concentrations, muscle supernatant protein levels were assessed using the Bradford method and diluted to equal concentrations prior to loading (pg/ml).

*RNA Isolation and Quantitative RT-PCR*

RNA was extracted from homogenized TA muscle using PureLink RNA Mini Kit (Cat. # 12183018A, Thermo Fisher Scientific, Waltham, MA, USA) following manufacturer's instructions. RNA was reverse transcribed into cDNA using a High-Capacity RNA to cDNA set (Cat. # 4368814, Thermo Fisher Scientific, Waltham, MA, USA). Taqman Fast Advanced MasterMix (Cat. # 4444556, Thermo Fisher Scientific, Waltham, MA, USA) was used for RT-PCR as per manufacturer's instructions.

**Figure Legends**

*Supplement 1*

CoO-TRIM reduces the frequency of small fibers and increases the number of large fibers following injury. Summary values comparing frequency (%) of **A**) small (CSA < 800 μm$^2$) and **B**) large (CSA > 2000 μm$^2$) fibers 8 dpi. Summary values comparing frequency (%) of **C**) small (CSA < 800 μm$^2$) and **D**) large (CSA > 2000 μm$^2$) fibers 14 dpi. Summary values are means ± SEM (n=8/group). Comparisons made by 2-Way ANOVA (condition x treatment), $p<0.05$ = significant

*Supplement 2*

Representative images of Healthy TA muscle cross-sections at 8- and 14 dpi. Laminin (white) identified basal laminae surrounding fibers, DAPI (blue) identified nuclei. Scale bars = 500 μm

*Supplement 3*

Anabolic signaling following CoO-TRIM treatmment. **3.1 and 3.2 A-D**) Representative immunoblots and mean densitometric data for phosphorylated, total, and ratio of phosphorylated to total protein abundance for Akt, mTOR, 4E-BP1, and p70s6k in whole muscle TA homogenate were normalized to total protein per lane, represented by the 40 kDA band from the total protein stain at 8- and 14 dpi. Summary values are means ± SEM (n=6/group). Comparisons made by 2-Way ANOVA (condition x treatment), $p<0.05$ = significant

*Supplement 4*

Proteolytic activity is unaffected by CoO-TRIM following injury. **A-D**) Representative immunoblots and mean densitometric data 8- and 14 dpi for αII-Spectrin at 250 kDA, and its cleavage products at 145 kDa (Calpain) and 120 kDA (Caspase-3) normalized to total protein per lane, represented by the 40 kDa band from the total protein stain. Summary values are means ± SEM (n=6/group). Comparisons made by 2-Way ANOVA (condition x treatment), $p<0.05$ = significant

*Supplement 5*

Representative merged images of TA muscle cross-sections at **5.1A and B**) 8 dpi and **5.2A and B**) 14 dpi. Laminin (white) identified basal laminae surrounding fibers, CD31 (green) identified endothelial cells. Scale bars = 200 μm

*Supplement 6*

Representative immunoblots and mean densitometric data for CD45 **A**) 8 dpi and **B**) 14 dpi. Summary values are means ± SEM (n=6/group). Comparisons made by 2-Way ANOVA (condition x treatment), $p<0.05$ = significant

*Supplement 7*

Chemokine and cytokine concentration *Top,* 8 dpi and *Bottom,* 14 dpi for **A**) IFNγ, **B**) IL-12p70, **C**) IL-2, **D**) IL-4, and **E**) IL-5. Summary values are means ± SEM (n=8/group). Comparisons made by 2-Way ANOVA (condition x treatment), $p<0.05$ = significant
