## Supplementary figures and images for "Skeletal Muscle Regeneration is Accelerated Following Injection of Time Release Ion Matrix in Injured Mice"

### Supplemental Figure 1

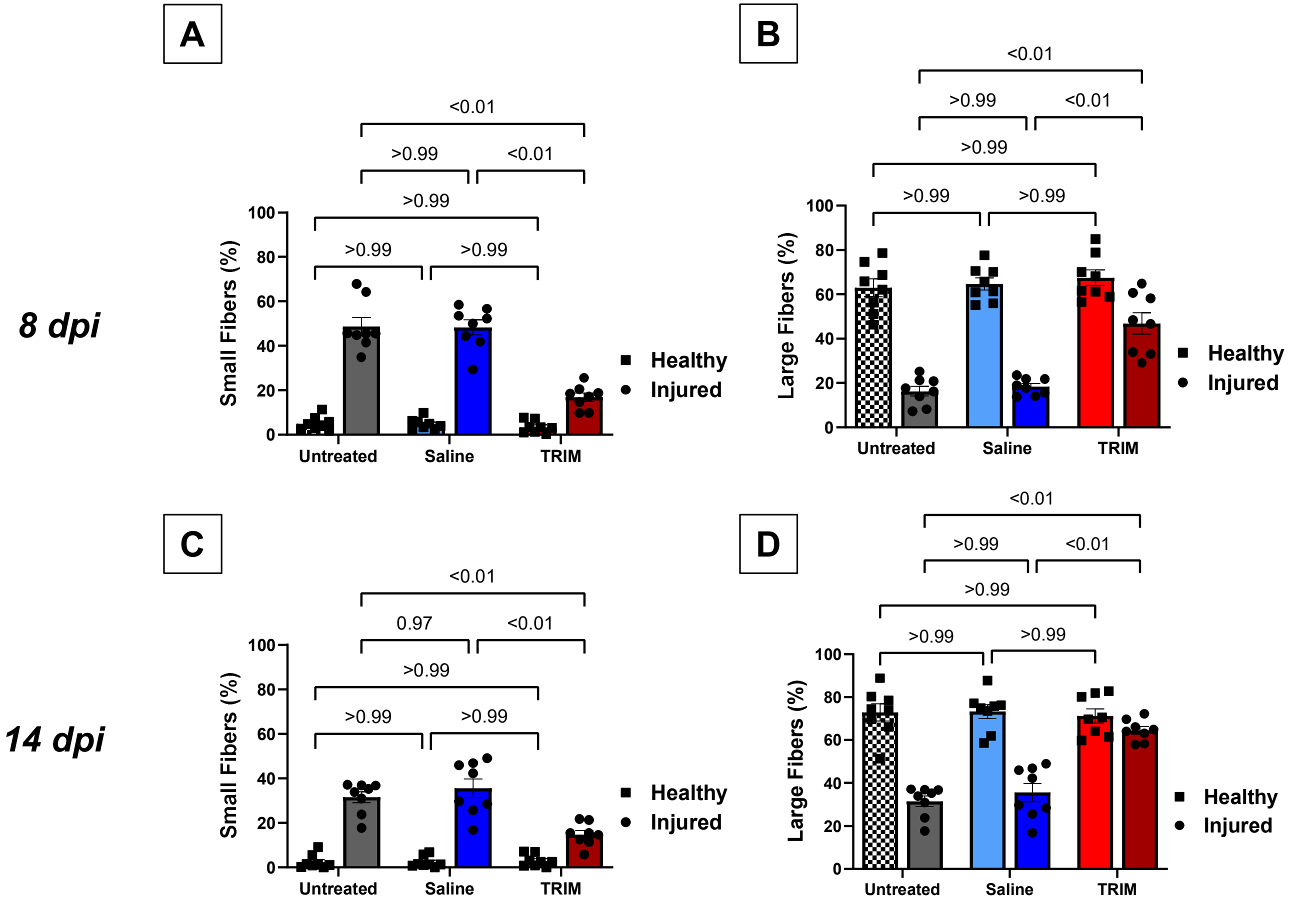

### Supplemental Figure 2

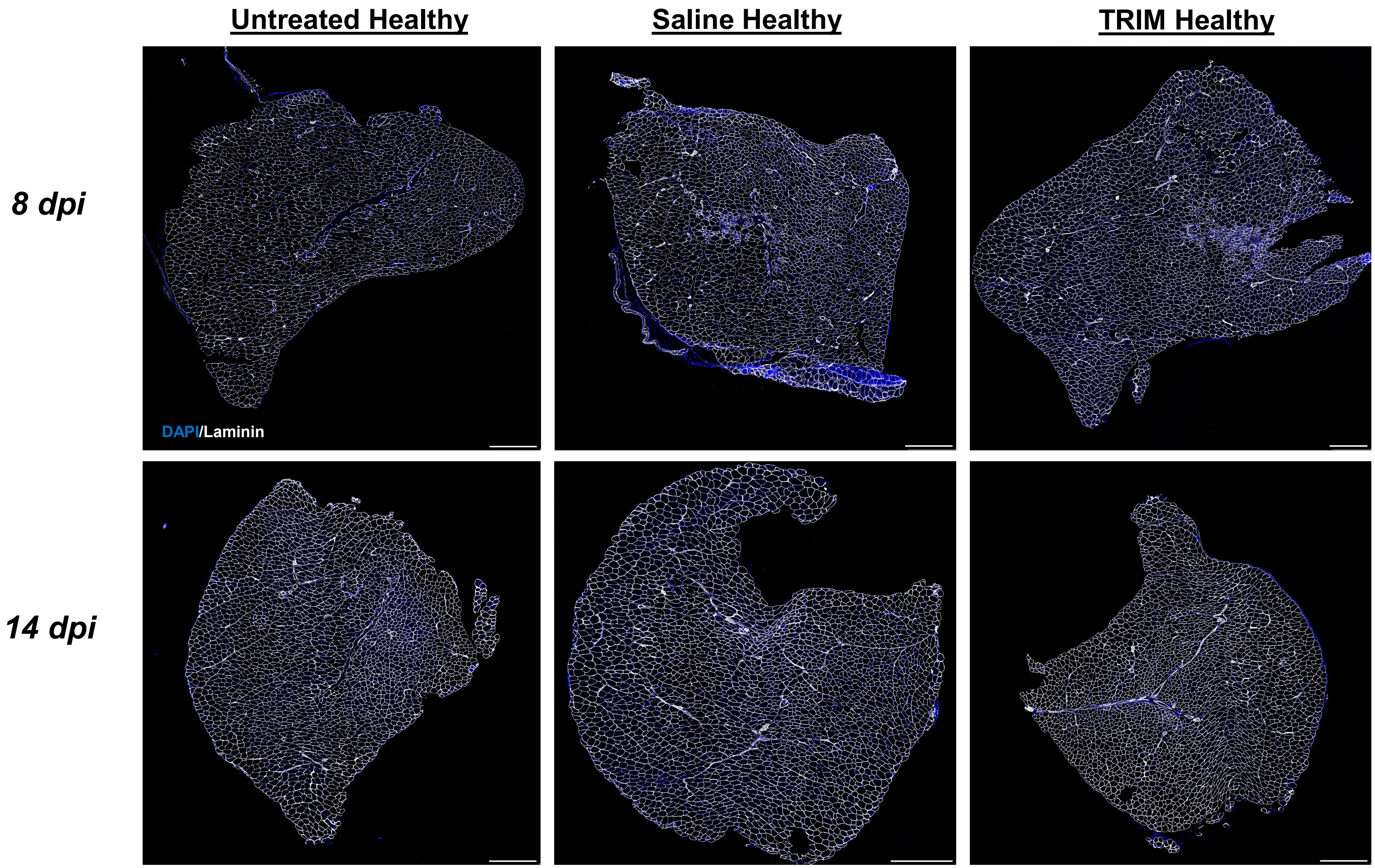

### Supplemental Figure 3.1

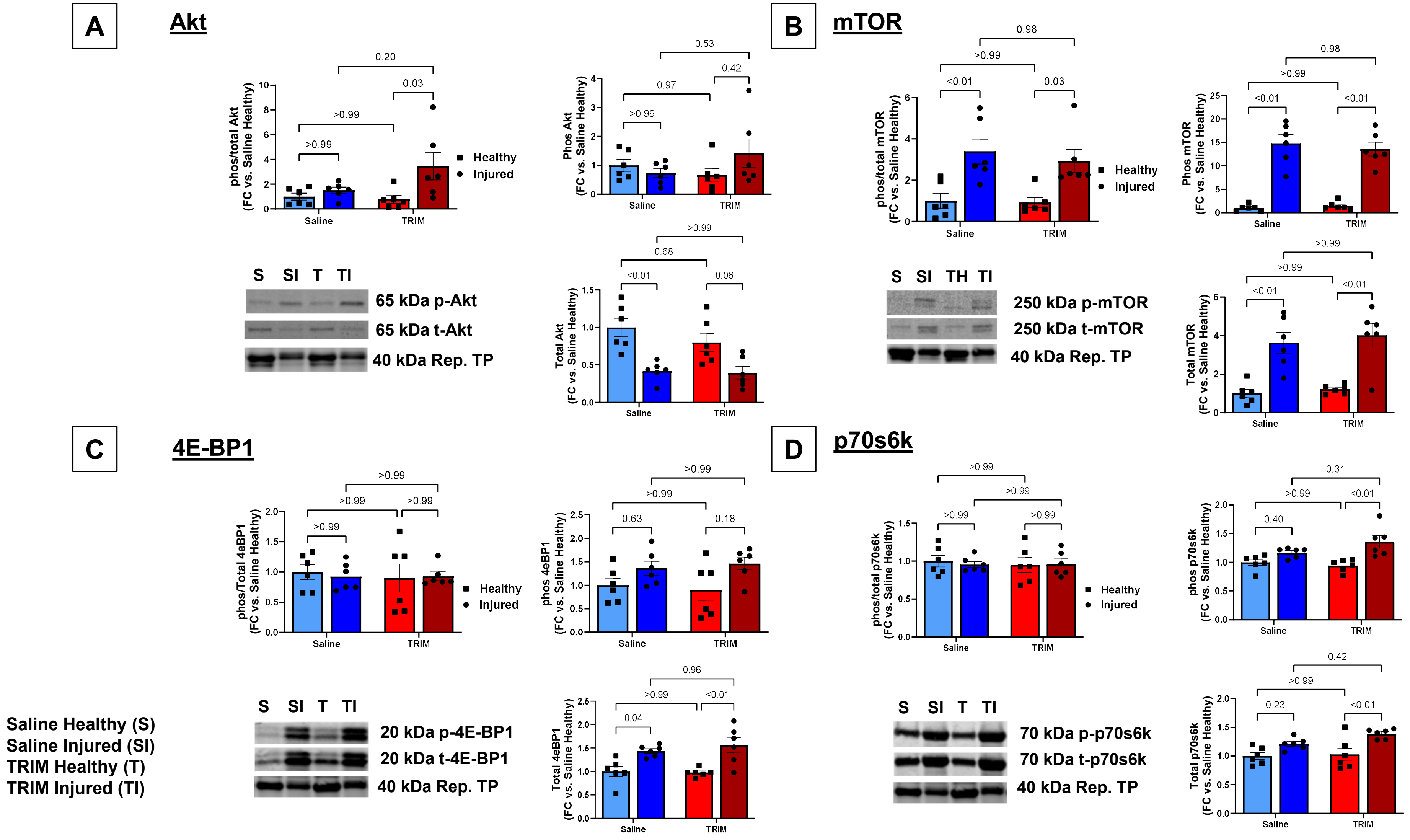

### Supplemental Figure 3.2

**S3.2**

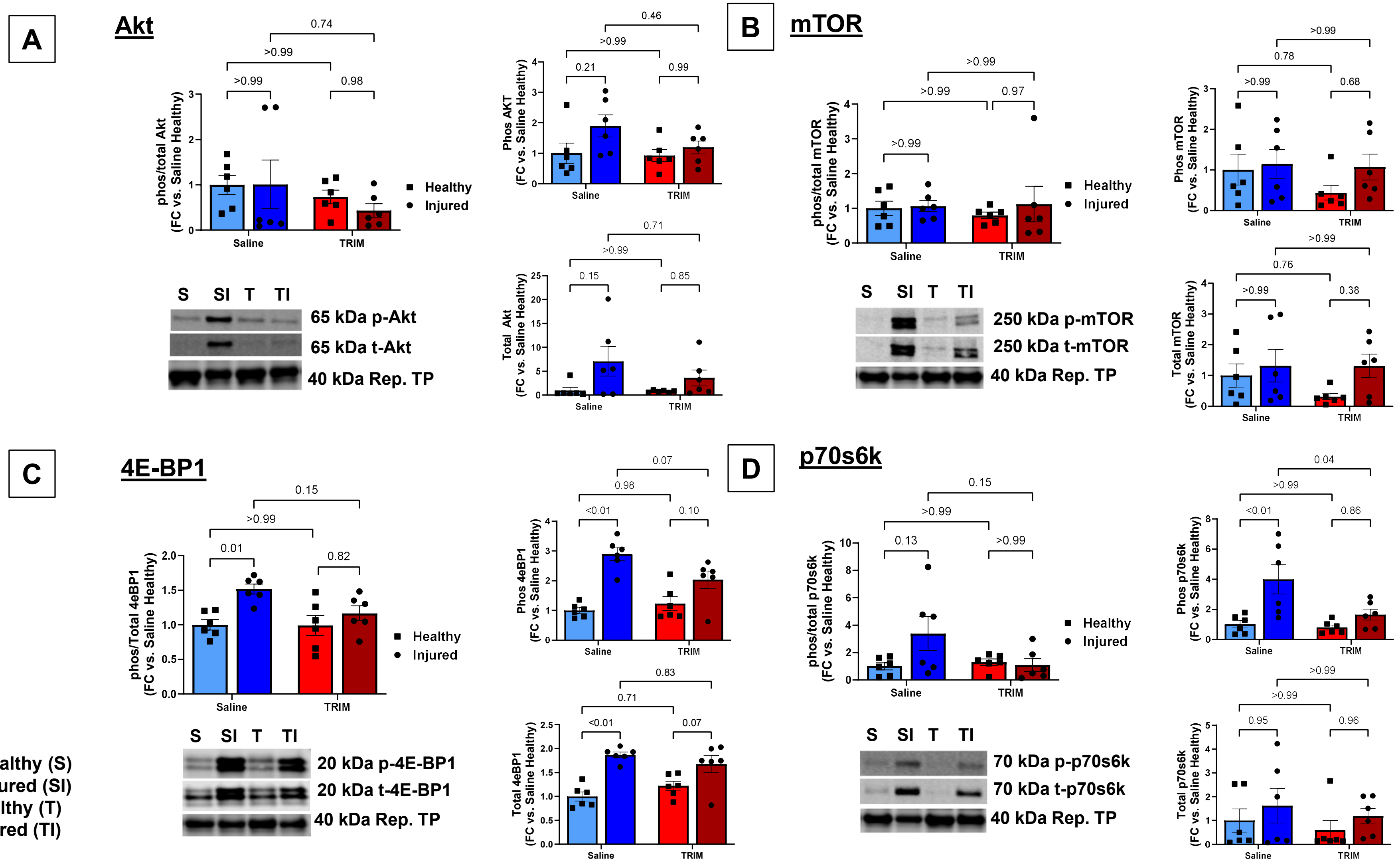


Saline Healthy (S)
Saline Injured (SI)
TRIM Healthy (T)
TRIM Injured (TI)

### Supplemental Figure 4

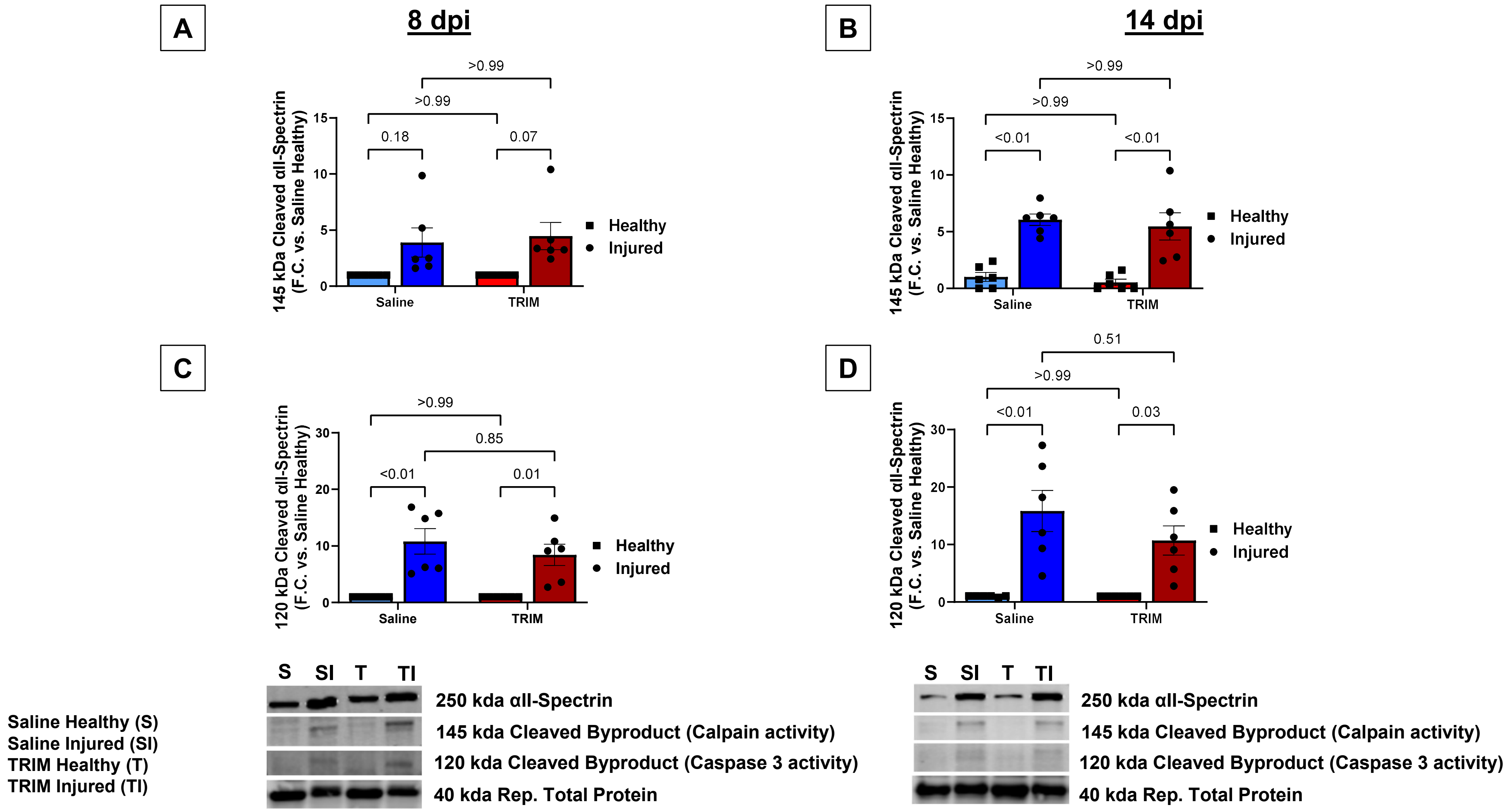

### Supplemental Figure 5.1

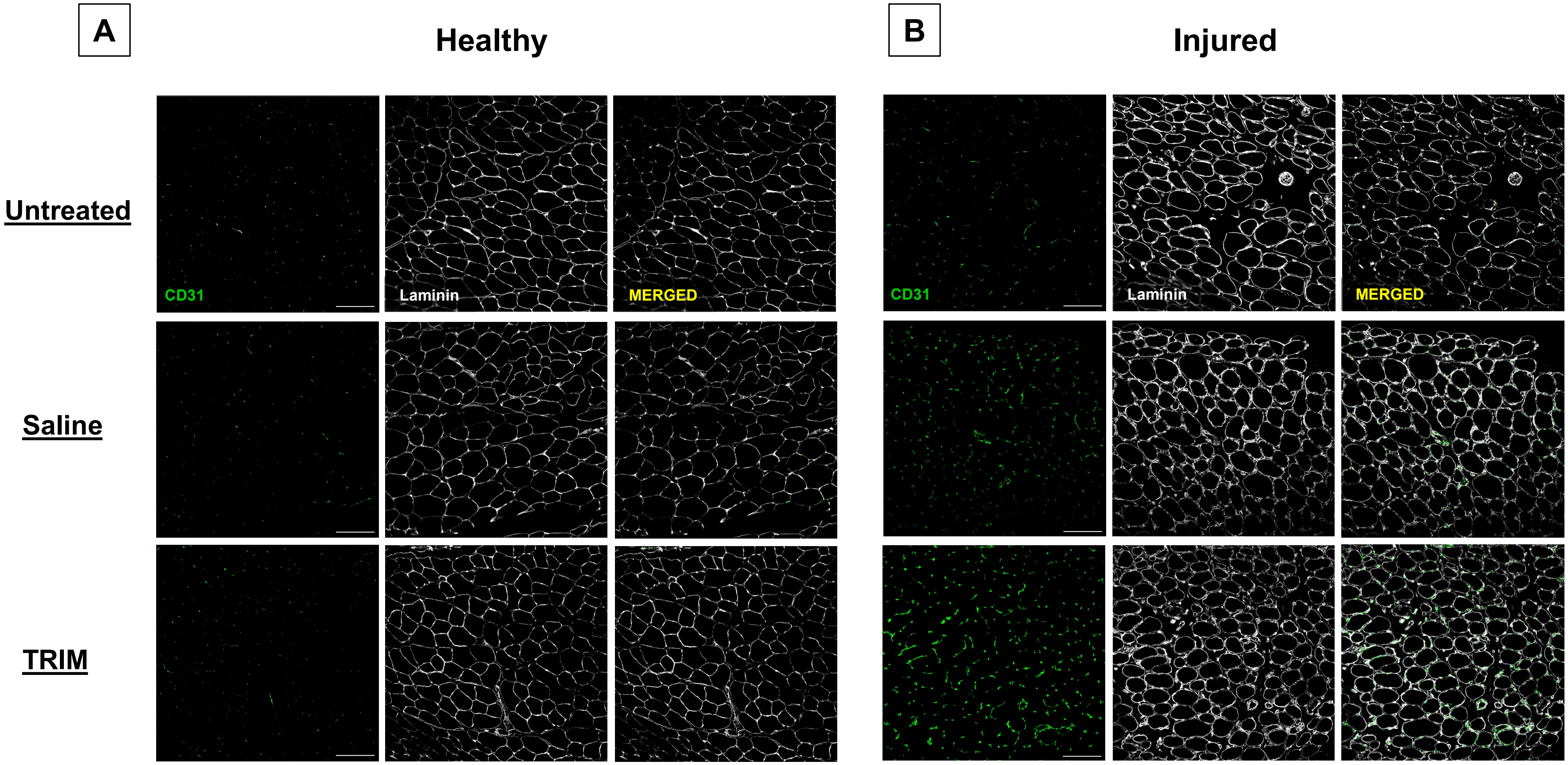

### Supplemental Figure 5.2

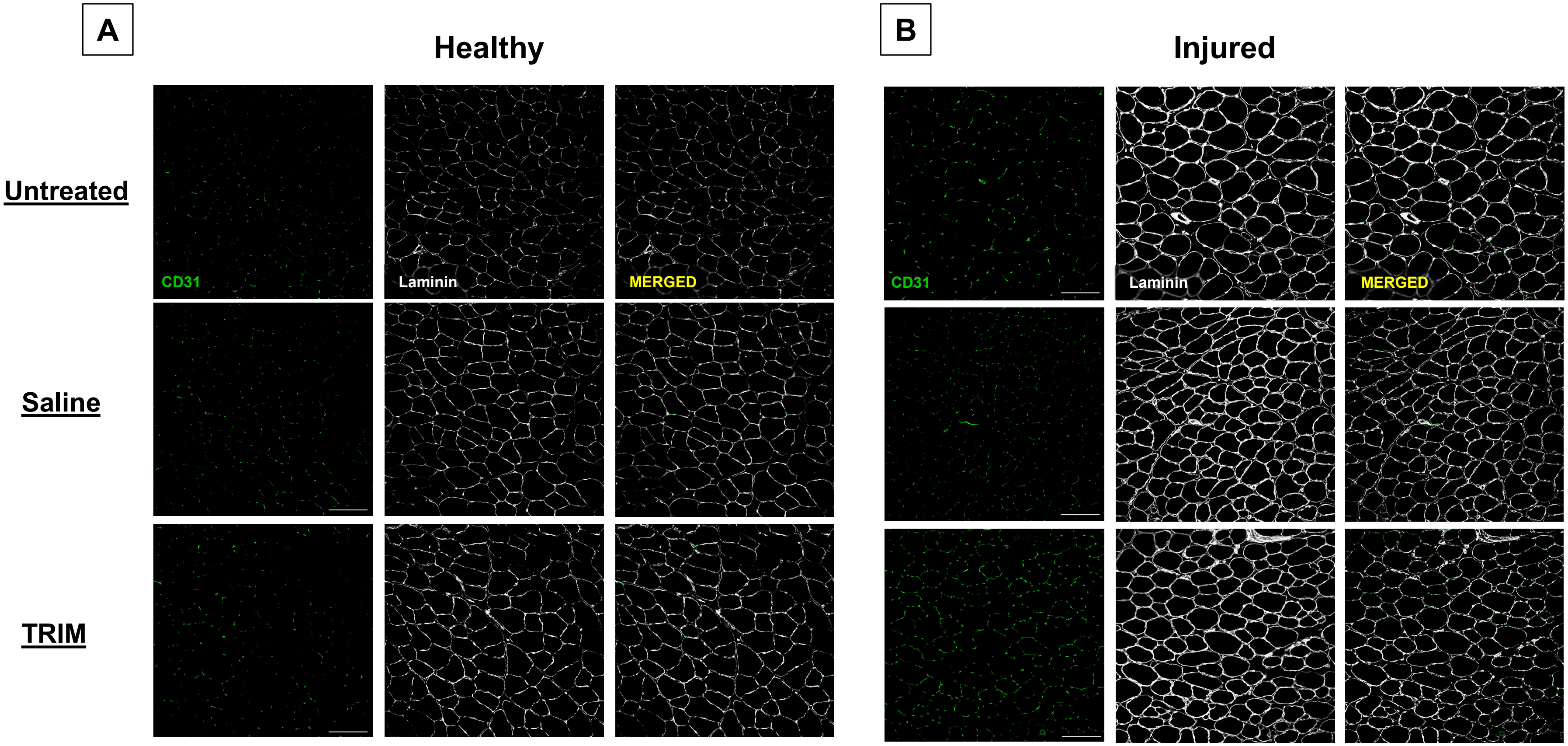

### Supplemental Figure 6

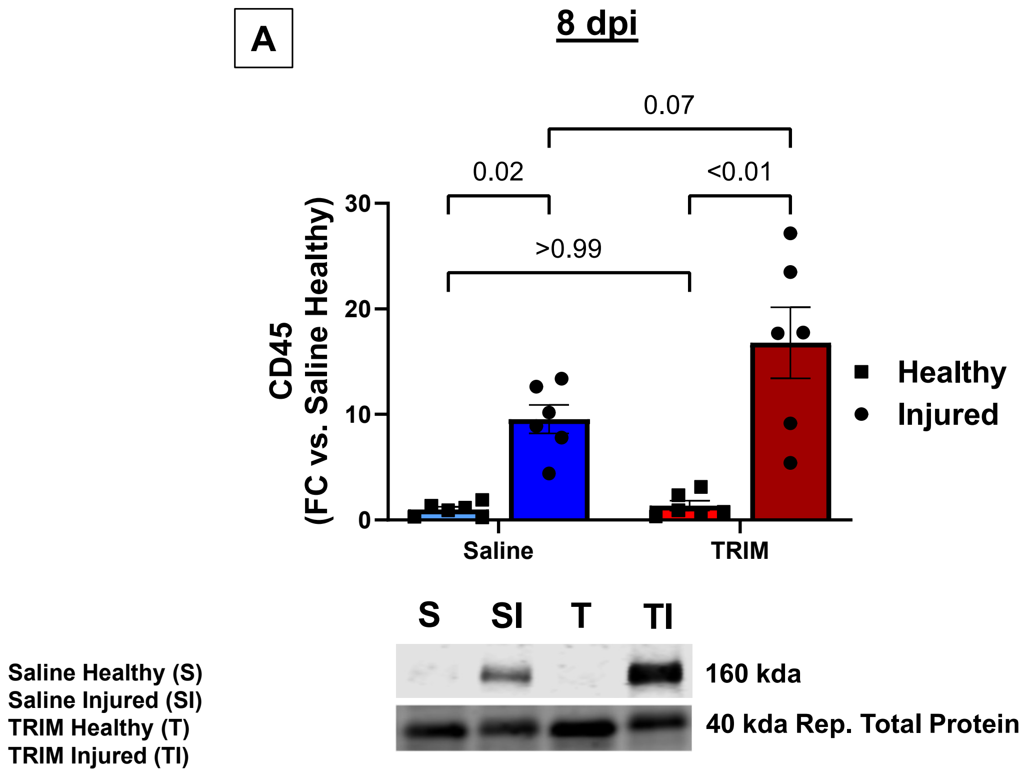

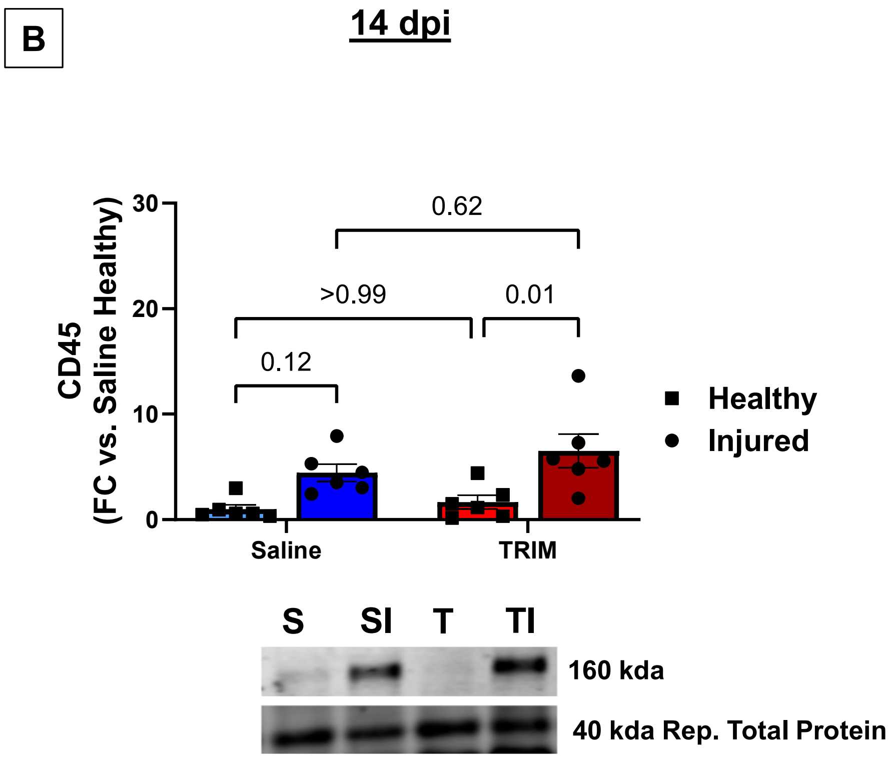

### Supplemental Figure 7

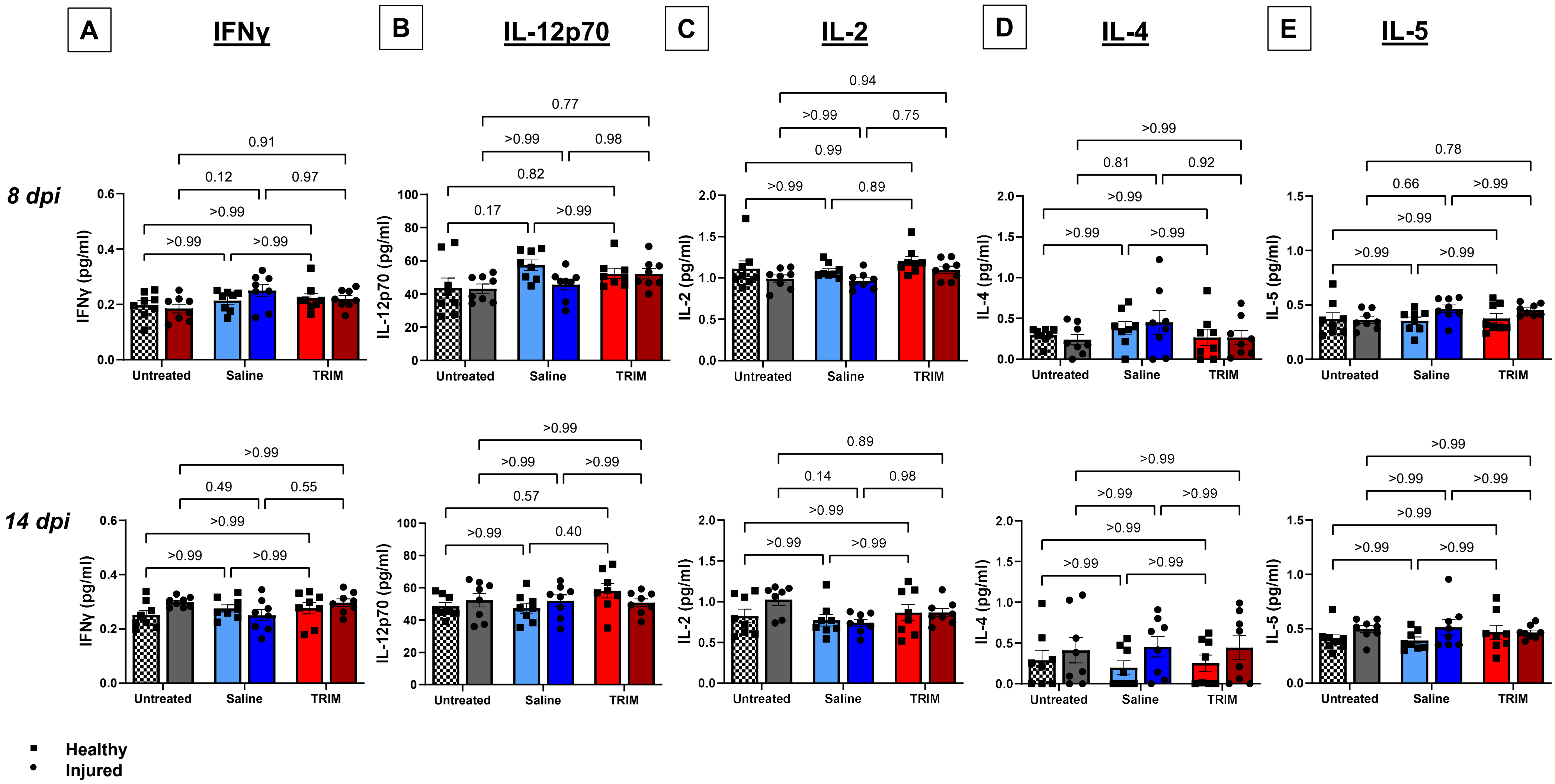
