## Supplementary material for "Skeletal Muscle Regeneration is Accelerated Following Injection of Time Release Ion Matrix in Injured Mice": Western Blot Supplement

### Immunoblot Membranes and Total Protein Stains

*Pax7*

**8 dpi**

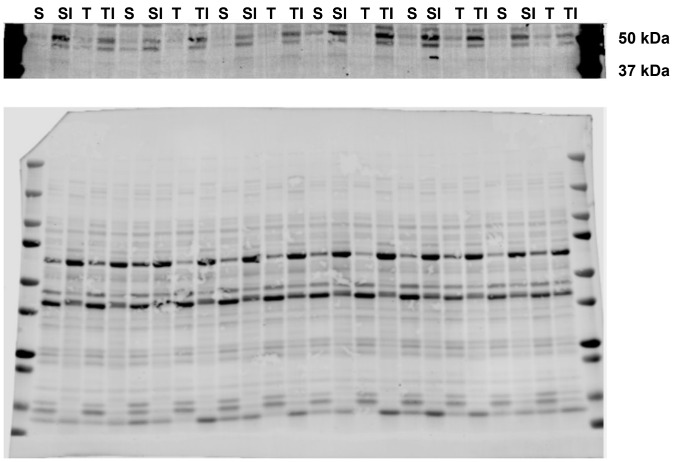


**14 dpi**

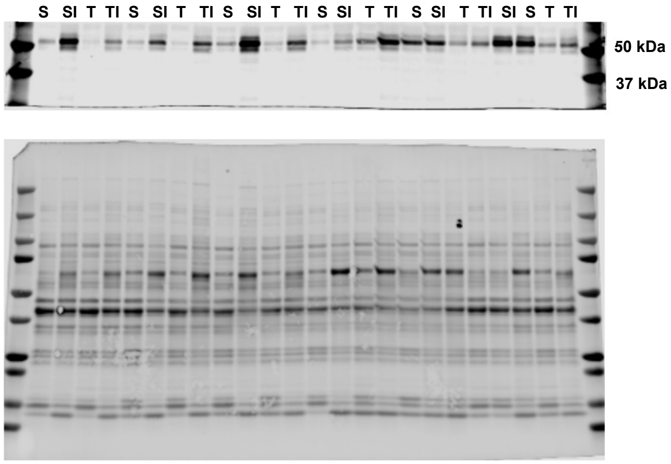


*MyoD*

**8 dpi**

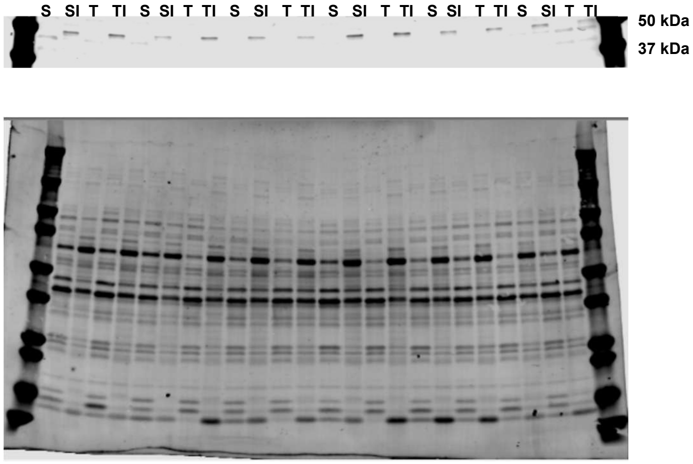


**14 dpi**

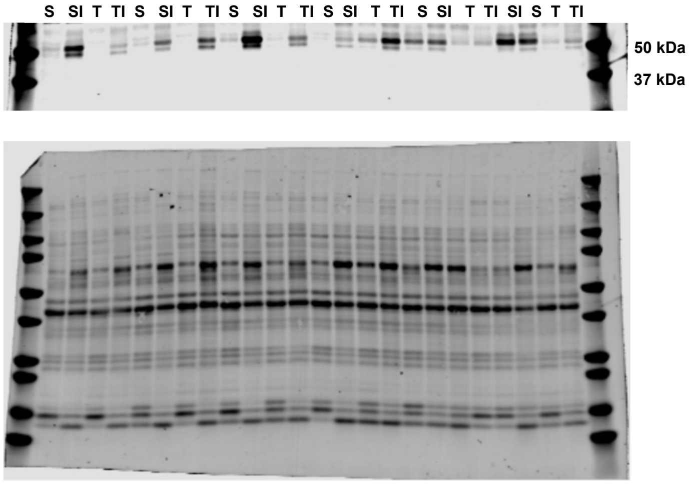

*Myogenin*

**8 dpi**

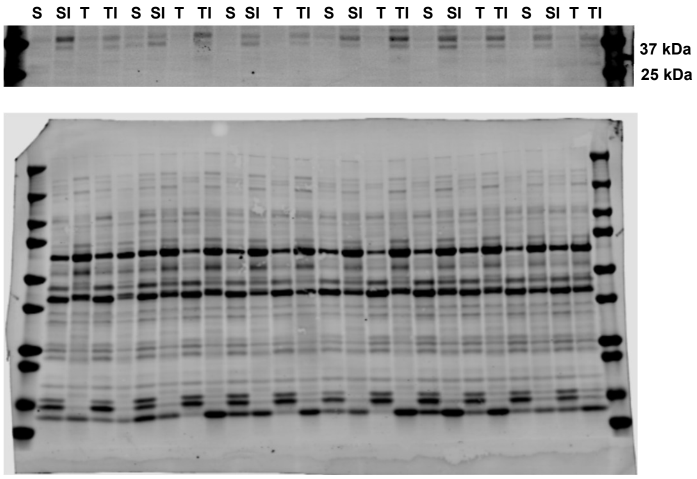


**14 dpi**

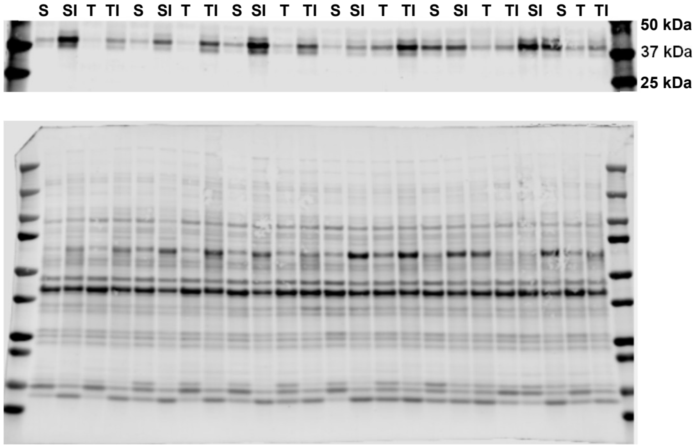


*CD45*

**8 dpi**

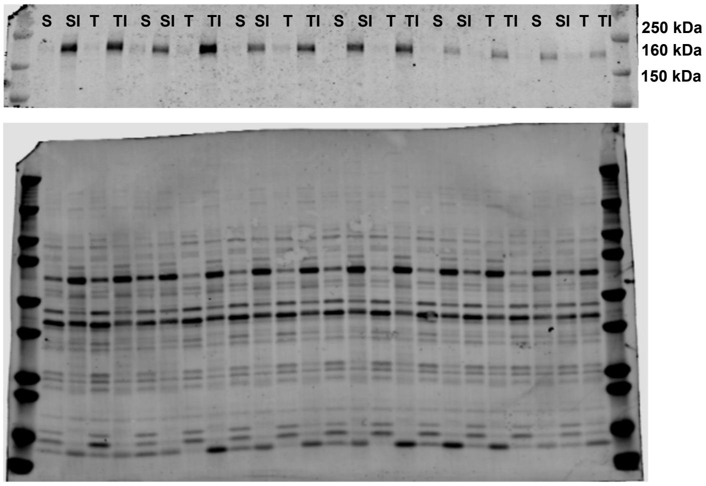


**14 dpi**

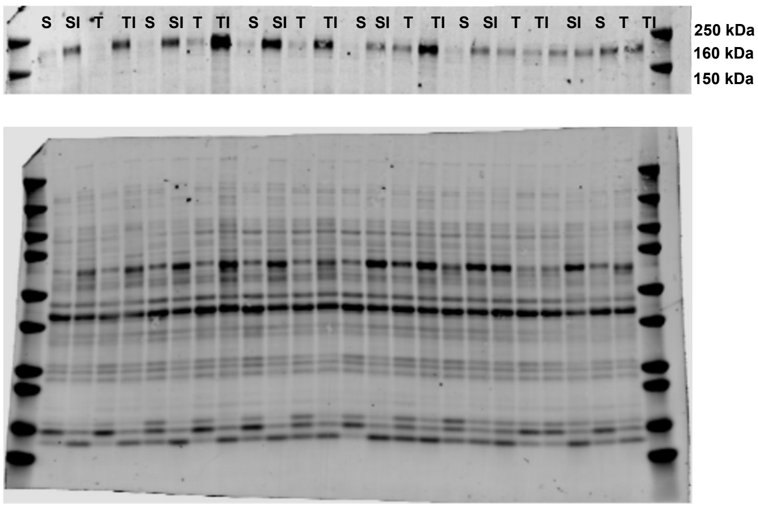

*Akt*

**8 dpi** | **14 dpi**

**Phosphorylated**

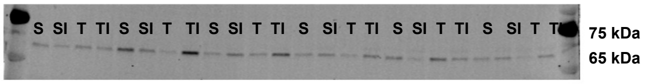


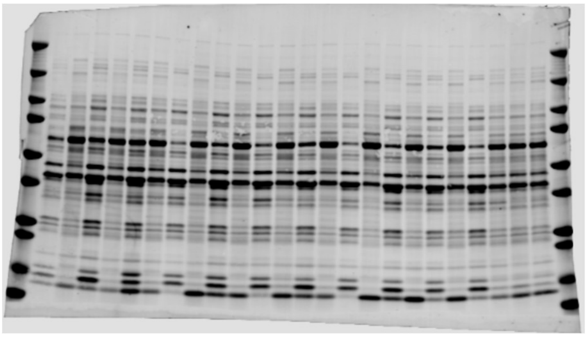

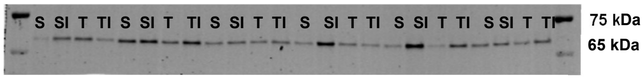


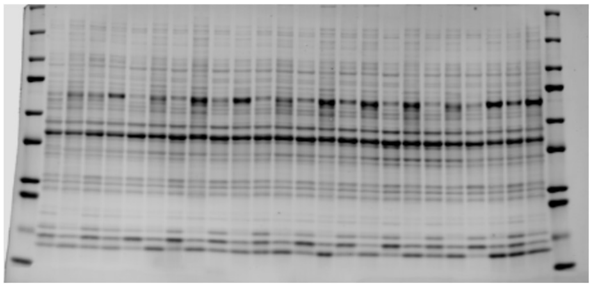

**Total**

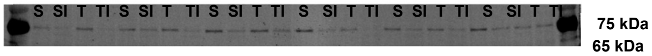


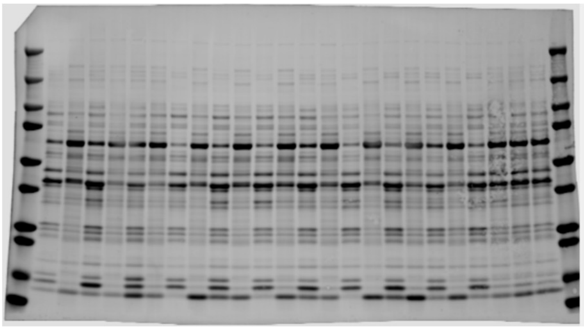

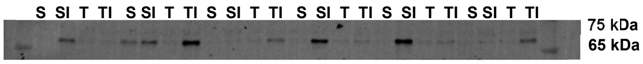


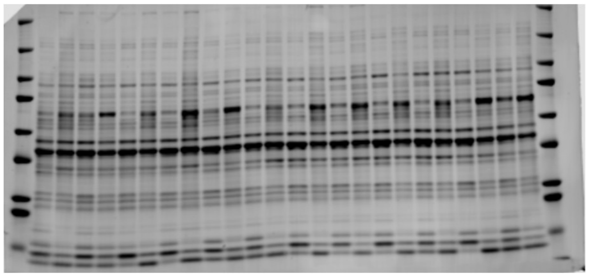

*mTOR*

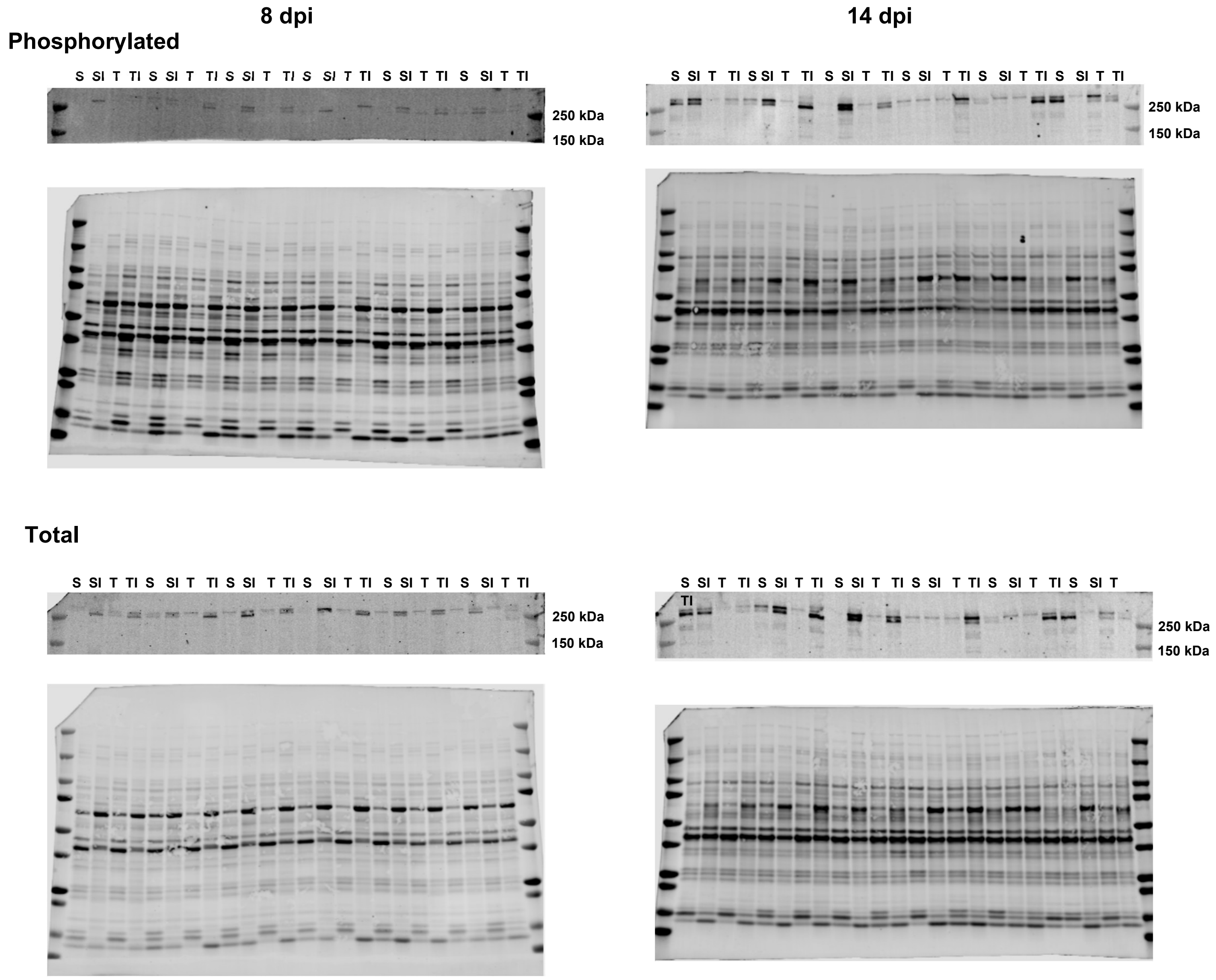
8 dpi
14 dpi
Phosphorylated
S SI T TI S SI T TI S SI T TI S SI T TI S SI T TI S SI T TI
250 kDa
150 kDa
S SI T TI S SI T TI S SI T TI S SI T TI S SI T TI S SI T TI
250 kDa
150 kDa
Total
S SI T TI S SI T TI S SI T TI S SI T TI S SI T TI S SI T TI
250 kDa
150 kDa
S SI T TI S SI T TI S SI T TI S SI T TI S SI T TI S SI T
TI
250 kDa
150 kDa

*4E-BP1*

**8 dpi** **14 dpi**

**Phosphorylated**

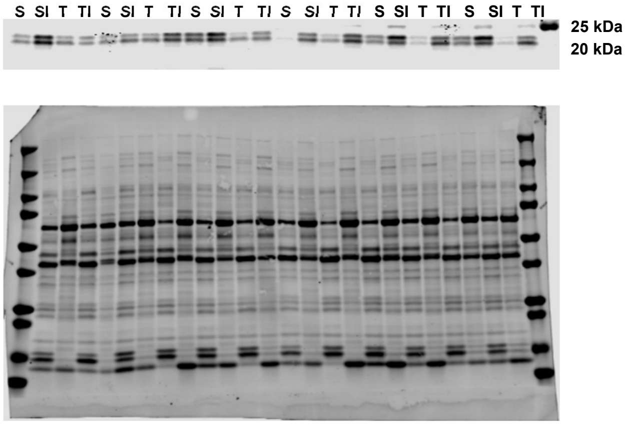


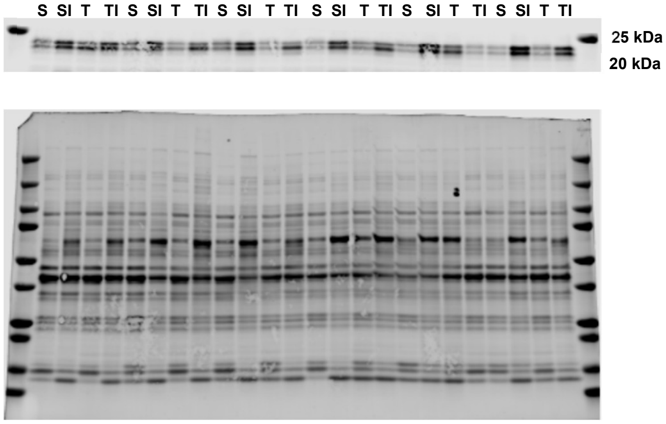


**Total**

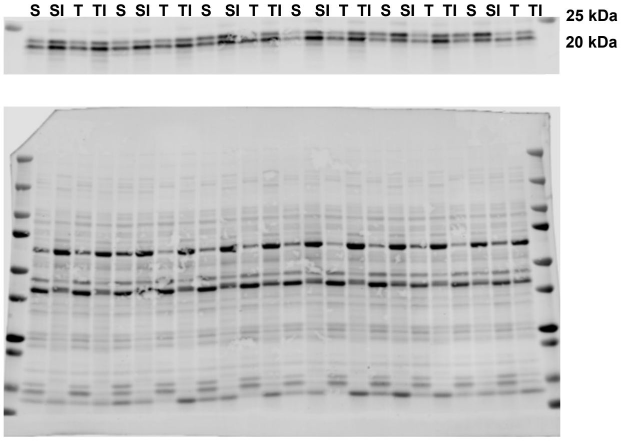


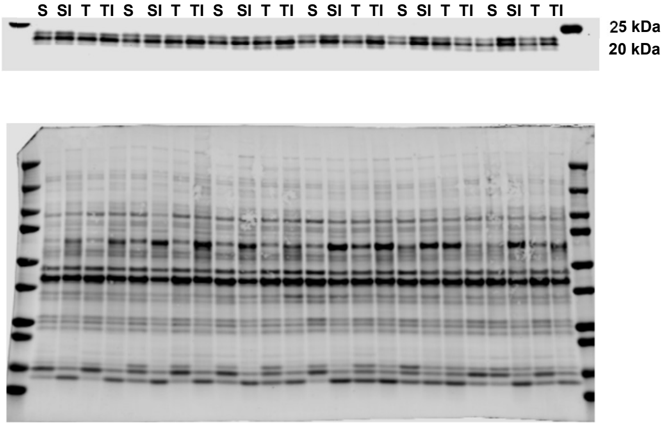

*p70s6k*

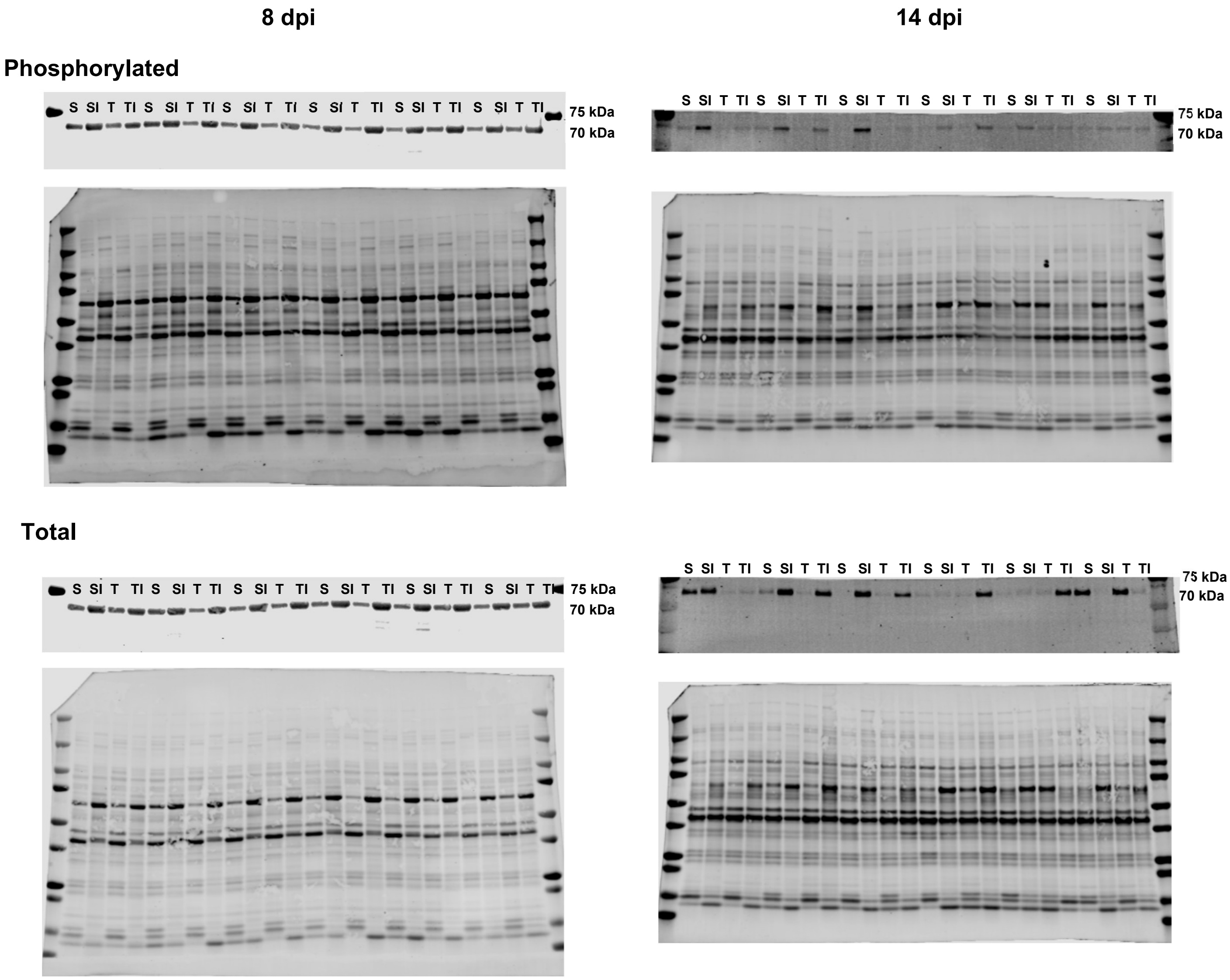
8 dpi
14 dpi
Phosphorylated
S SI T TI S SI T TI S SI T TI S SI T TI S SI T TI S SI T TI
75 kDa
70 kDa
S SI T TI S SI T TI S SI T TI S SI T TI S SI T TI S SI T TI
75 kDa
70 kDa
Total
S SI T TI S SI T TI S SI T TI S SI T TI S SI T TI S SI T TI
75 kDa
70 kDa
S SI T TI S SI T TI S SI T TI S SI T TI S SI T TI S SI T TI
75 kDa
70 kDa

*αII-Spectrin*

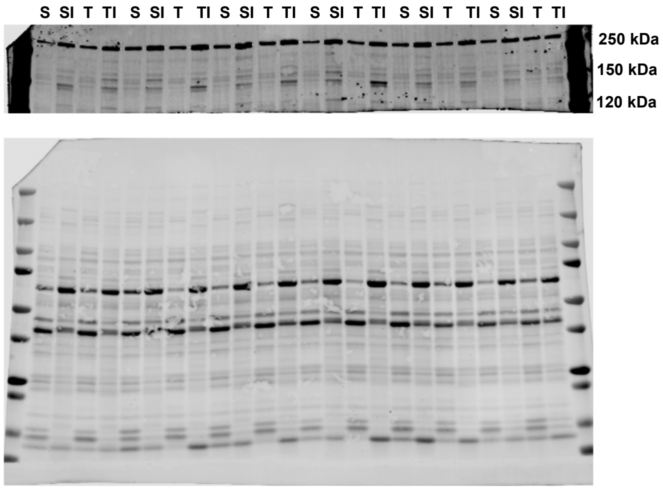


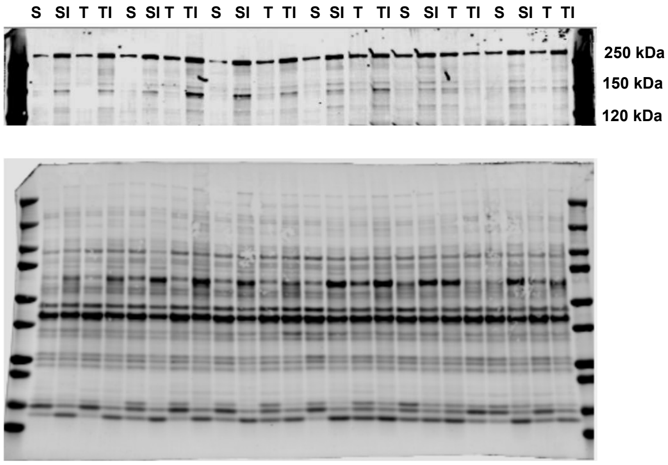
